# Beyond the Focus of Expansion: Retinal curl as a functional signal for heading estimation

**DOI:** 10.64898/2026.01.21.700857

**Authors:** Kontessa I. Zorpala, Joan López-Moliner

## Abstract

Prevailing models aiming at explaining heading assume that humans need to recover the Focus of Expansion (FoE) while accounting for eye-movement-induced rotation. We propose an alternative: the visual system utilizes mean retinal curl from fixations as a surrogate signal for heading, rendering the explicit recovery of the FoE unnecessary. Stationary participants viewed simulated walking paths on a large screen while fixating on points on the projected ground texture at varying eccentricities —a natural behavior inducing sustained retinal curl. Participants continuously reported perceived heading in 3D scene coordinates. To isolate the role of retinal curl, we employed a real-time manipulation that kept translational flow constant while the foveal curl component was either unaltered, cancelled, or overcancelled. Under natural conditions (unaltered), participants exhibited systematic heading biases opposite the direction of gaze. Crucially, these biases vanished when we cancelled the expected curl, and flipped when we overcancelled it, identifying retinal curl as the specific driver of perceptual bias. We modeled these results using a simple feedback controller and a ring-attractor neural network featuring gaze-contingent inhibition and a ‘straight-ahead’ prior. These findings suggest the brain exploits the geometry of gaze stabilization to simplify navigation, treating retinal curl as a functional signal rather than noise to be filtered.

## Introduction

Humans and many vertebrates rely on stabilizing gaze on regions of interest in the world to achieve accurate perception. This fixation strategy, combined with the continuous eye, head, and body movements that accompany natural behavior, generates highly structured patterns of motion on the retina that can, in principle, be exploited to control locomotion. A particularly simple and influential case arises when we move approximately in the same direction as we are looking: in this situation the focus of expansion (FoE), the point in the optic flow from which motion vectors appear to radiate, directly specifies the heading or trajectory direction (Gibson, 1950). The idea that the visual system uses the FoE to recover heading has subsequently dominated both theoretical and empirical work, including the interpretation of neural activity in primate medial superior temporal (MST) area, where neurons exhibit FoE-like tuning (Bremmer et al., 2017; Britten, 2008; Duffy and Wurtz, 1991; Kaminiarz et al., 2014), as well as psychophysical findings showing that humans can make highly accurate heading judgements from radial optic flow (e.g Warren and Hannon (1990); Warren and Hannon (1988); Warren et al. (1988); van den Berg (1992); Li and Warren (2000)).

However, rotating the eyes or head to look at an off-path object adds a rotational component to the retinal flow. This combined motion distorts the pure expansion pattern, shifting the FoE so it no longer aligns with the true heading. Despite this distortion, observers can generally maintain accurate heading judgments during both active eye movements (Banks et al., 1996; Royden et al., 1992) and simulated rotations—where the rotational visual component is artificially rendered on a screen without the observer actually moving their eyes—provided sufficient 3D depth is visible (Cutting, 1986; Grigo and Lappe, 1999; Li and Warren, 2002; Warren and Hannon, 1988). To explain how humans distinguish where they are heading from where they are looking, classic models suggest the brain decomposes the flow or subtracts the rotation to recover the pure FoE. This de-rotation is achieved either by exploiting visual cues alone (Beintema et al., 2004; Cutting et al., 1992; Heeger and Jepson, 1992; Lappe and Rauschecker, 1993; Longuet-Higgins and Prazdny, 1980; Perrone and Stone, 1994), or by integrating extra-retinal signals (Beintema and van den Berg, 1998; Koenderink and van Doorn, 1981; Lappe, 1998; Royden et al., 1994; van den Berg and Beintema, 2000). Yet, this compensation is not perfect: participants often make systematic heading errors during these simulated rotations devoid of extra-retinal cues (Banks et al., 1996), particularly when depth is absent (such as when viewing a flat, single fronto-parallel wall) (Grigo and Lappe, 1999; Rieger and Toet, 1985) or when the tracking target moves independently of the ground plane (Royden et al., 1992).

The fact that recovering self-motion from rotational flow requires some type of parallax (e.g. Rieger and Toet (1985); Stone and Perrone (1997); Grigo and Lappe (1999)), depth order (van den Berg and Brenner, 1994) or rigid 3D structure to guide heading (Cutting, 1986; Li and Warren, 2000; Warren, 1998) and steering (Li and Warren, 2002; Wann and Swapp, 2000) directly challenges visual decomposition models. If the brain simply subtracted rotation from translation, heading could theoretically be recovered even from flat, fronto-parallel planes. The systematic errors observed when depth is absent—unless extra-retinal cues are available— suggest the brain bypasses global de-rotation. Pure visual decomposition should function regardless of 3D depth or whether the rotation stems from an active eccentric fixation.

Indeed, behavioral evidence shows that humans successfully recover heading specifically when this rotational flow arises from tracking an eccentric, grounded target (Perrone and Stone, 1994; van den Berg, 1993; Warren and Hannon, 1988). In natural navigation, this type of continuous gaze stabilization is ubiquitous (Matthis et al., 2018). Instead of creating a nuisance artifact that must be subtracted, this gaze behavior generates a useful, depth-dependent rotational component in the form of structured spiral gradients—the retinal curl. In real-world locomotion, foveal curl provides a more reliable gaze-relative signal than the highly variable, head-centered FoE (Matthis et al., 2022). Crucially, this curl extends beyond estimating an instantaneous heading to specify a “future path” (Wann and Swapp, 2000; see Li, 2025 for a recent review), offering predictive information that is highly advantageous for active steering. The use of curl aligns with reinterpreting neurophysiological evidence: rather than utilizing segregated expansion and rotation channels, MSTd neurons would be tuned to a continuum of spiral motion patterns (Graziano et al., 1994; Layton and Browning, 2014). Such tuning suggests the visual system actively exploits the rotational gradients inherent to active fixation strategies (Angelaki and Hess, 2005; Calow and Lappe, 2008; Glennerster et al., 2001).

Here, we provide empirical evidence that the visual system utilizes retinal curl as a primary control variable for locomotor navigation. Participants reported their perceived trajectory in world-centered (3D scene) coordinates during simulated walking through a naturalistic optic flow field (see Figure 1A). This paradigm revealed a robust, systematic bias directed strictly opposite to the direction of gaze: fixating an eccentric ground target to the left induces a rightward shift in perceived heading, and vice versa. This effect, which requires sustained exposure and differs qualitatively from classic simulated-rotation errors (Supplementary Videos S1 and S2), is precisely predicted by gaze-centered flow geometry. Crucially, experimentally cancelling the retinal curl eliminated the bias entirely, confirming its causal role. To mechanistically ground these findings, we show that a simple feedback controller driven by mean image curl faithfully reproduces human steering behavior across diverse path conditions. Finally, we provide a neurally plausible implementation illustrating how parietal circuits might transform local foveal curl into global heading estimates via recurrent dynamics and the competitive integration of sensory evidence and spatial priors. Consequently, we propose a redefined role for extra-retinal information. Rather than simply “de-rotating” the flow field, eye position signals serve to scale and interpret retinal curl. The visual system would exploit the magnitude of foveal curl directly, transforming this structured spiral gradient into calibrated steering commands without uncovering a hidden FoE.

**Figure 1:**
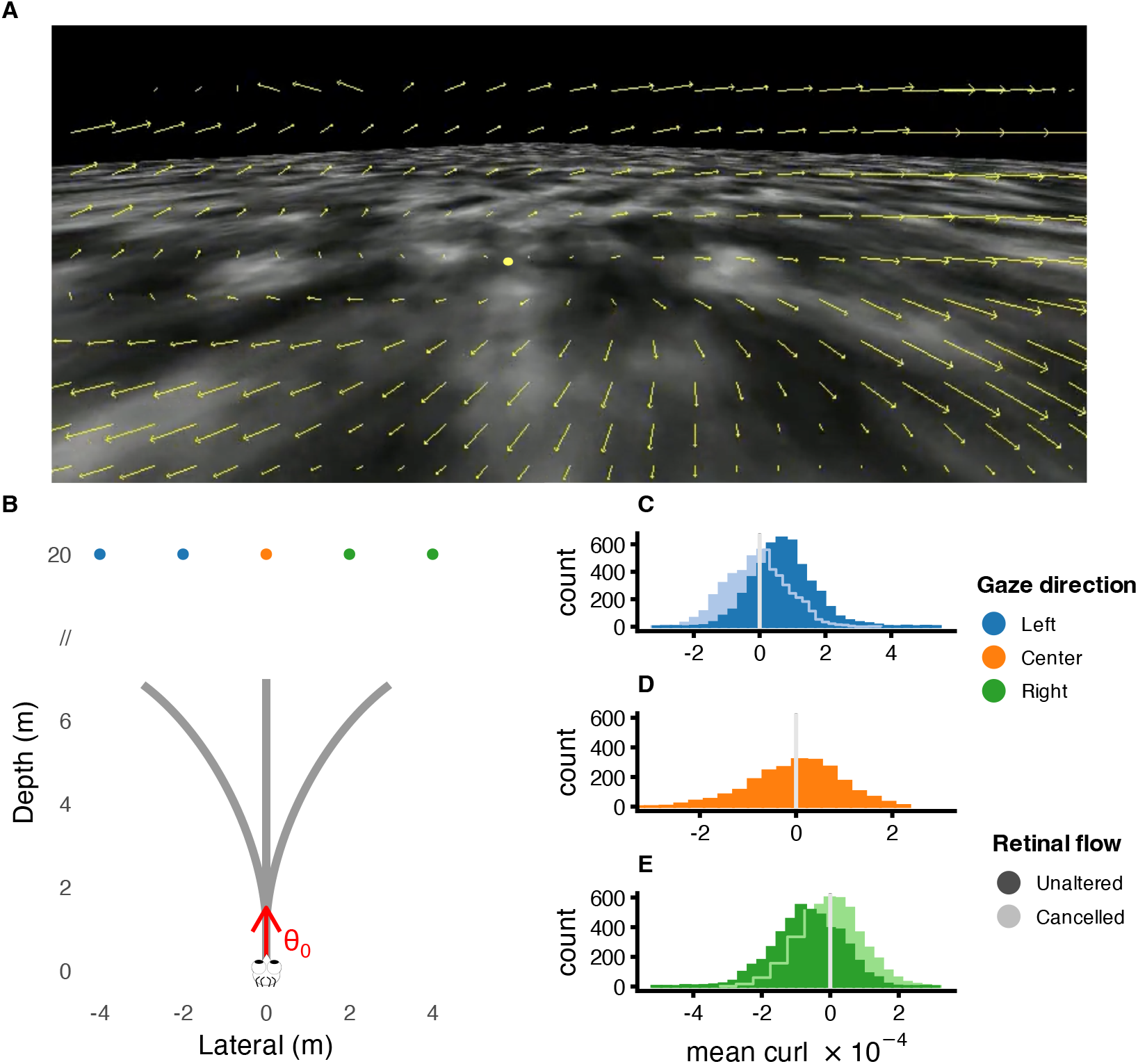
Ground texture, trajectories and retinal curl distributions across conditions. (A) Snapshot of the ground texture based on simplex noise. Yellow lines indicate optic flow vectors computed using the Farnebäck algorithm. A clear rotational component (curl) is visible, consistent with the observer looking at a point (yellow dot) located to the left of the simulated path. The Focus of Expansion is shifted in the direction of the gaze. (B) Schematic of the experimental trajectories. Position (0, 0) represents the starting point of simulated locomotion. The red arrow indicates the initial heading (*θ*_0_), which was aligned longitudinally with the 3D scene (world coordinates). Participants were instructed to report their perceived heading within this world-centered reference frame. The five colored dots mark the fixation points in world coordinates at the beginning of each trial (20 m ahead of the observer). (C–E) Distributions of mean retinal curl across trials for left-, center-, and right-gaze conditions, respectively. Dark filled bars indicate the unaltered curl condition, while lighter bars and outlined steps represent the cancelled curl condition. The vertical grey line denotes zero curl.

## Results

Observers viewed simulated forward locomotion along different paths over a ground plane (Figure 1B). During each trial, they maintained continuous fixation on a specific ground-embedded target, which was either aligned with their path (straight ahead) or laterally displaced (2 m or 4 m to the left or right); generating different mean curl distributions (see Figure 1C-E). While keeping this stabilized gaze, participants continuously reported their perceived heading in world coordinates, allowing us to reconstruct their full perceived trajectory over time.

We first assessed the quality of fixation to ensure gaze stability. Across subjects, the median trial-averaged deviation from the fixation target ranged from 1.02^°^ to 1.55^°^. Gaze-to-target distance exceeded 3^°^ in only 0.46% of trials; these were excluded from further analysis. In addition to spatial accuracy, we calculated the retinal slip (the velocity of the target image on the retina). Median horizontal slip ranged from 0.12 to 0.27^°^/*s* and vertical slip from 0.12 to 0.23^°^/*s*, confirming that gaze was effectively stabilized throughout the simulated locomotion.

Figure 2 shows the instantaneous perceived heading across different gaze and path conditions where retinal flow was unaltered. Paths were reconstructed in world (3D scene) coordinates by integrating the raw responses (see *Computation of Perceived Path* in the Methods section and Figure 2 Suppl. Figure 1 for an example of raw responses). The thin traces show this pattern for individual observers, and the thick coloured traces show the group means. Figure 2 illustrates a clear and systematic lateral bias in perceived heading as a function of the inital gaze direction. When observer-simulated motion is straight but fixated an eccentric point on the ground (central vertical panels, blue and green lines), their reported instantaneous heading consistently shifted away from the fixation point: leftward fixation led to rightward heading reports, and rightward fixation produced the opposite pattern. The perceived final lateral displacement is significantly different from 0 (straight ahead, t(11)=5.172, p<0.001) and the magnitude of the bias was not different between sides (F(1,35)<1, p=0.4). This effect was present both when gaze was displaced by 2 m (top row) and 4 m (bottom row), and the effect scales with the lateral distance of the fixation point from the trajectory path (the final lateral distance between 2 m and 4 m condition was marginally significant, F(1,35)=3.49, p=0.07). As we shall see below, the magnitude of the bias is well accounted for by the present mean curl.

**Figure 2:**
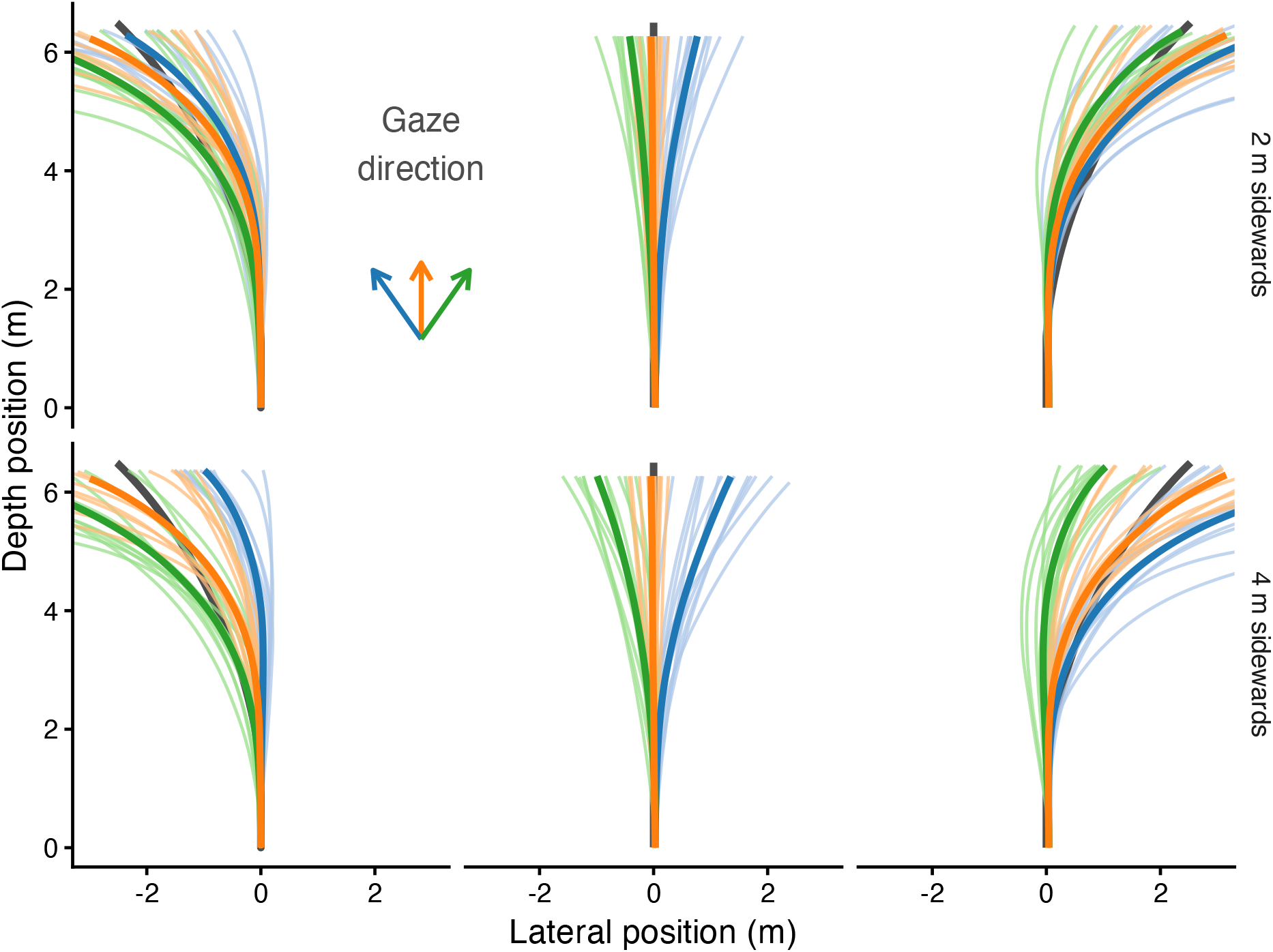
Perceived heading. Reported instant directions are plotted when fixating eccentric points on the ground. Rows: fixation 2 m (top) and 4 m (bottom) to the side. Columns: different physical path conditions (centre column = straight path). Colours code for initial gaze direction (left / centre / right). Thick coloured lines denote the mean across observers. Thin coloured lines denote individual observers. Dark grey denotes physical paths. Axes show lateral position (x-axis) versus depth position (y-axis)

Note that when both simulated motion and gaze are straight ahead (orange traces), there is essentially no bias: path is perceived straight. This is consistent with curl staying near zero throughout the trial (see Figure 1D). In this case, heading and gaze remain aligned and perceived trajectory stays close to the physical straight path.

Although the bias is more easily interpretable in the straight-ahead condition, systematic effects due to gaze direction are also present in curved paths. Perceived curvature was systematically *underestimated* when gaze was directed into the direction of the physical curve: for leftward curves, this underestimation is most evident when gaze is to the left (blue traces), and for rightward curves, when gaze is to the right (green traces).

This follows directly from the geometry of gaze-stabilization. As the observer proceeds along a curved path while looking toward the inside of the turn, the heading (*θ*_*t*_) and gaze directions (*ψ*_*t*_) become progressively aligned. This alignment causes the magnitude of retinal curl to naturally diminish over time. In our proposed controller (see Methods: control model), the visual system treats this curl as an error signal (*ϕ*_*t*_ = *ψ*_*t*_ − *θ*_*t*_) that drives the required turning rate: the yaw command to change heading 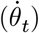 scales with this error (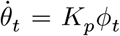, see Equation 6). Therefore, as the curl signal ‘dies out’ due to the alignment of gaze and heading, the model predicts a corresponding reduction in the perceived turning command—resulting in a progressive flattening of the perceived path. This is exactly the pattern observed empirically, and it is symmetric for left and right curves. By contrast, when gaze is directed opposite to the direction of curvature, the gaze and heading do not align —*ϕ* does not collapse toward zero—, the curl remains sustained, and no underestimation of curvature is expected or observed.

### Effects of flow manipulation

To establish a direct causal link between retinal curl and the observed heading biases, we introduced specific manipulations to the optic flow field in interleaved trials. By artificially cancelling or over-cancelling the natural rotational flow generated during eccentric fixation, we tested whether the participants’ perceived trajectories would predictably flatten or reverse in agreement with the exposed curl.

Figure 3 shows the reported instant heading for the different flow manipulations. We also show the unaltered condition (green) again for the sake of comparison. Importantly, when curl is cancelled (cyan), these biases essentially disappear – the perceived path remains much closer to the physical trajectory across gaze directions. Note that the central row (straight-ahead simulated motion) provides the clearest demonstration: the unaltered curl shows the previously reported lateral biases, but when curl is cancelled (counteracted) the perceived trajectory becomes near-veridical. This confirms that the mean curl signal itself – not gaze eccentricity per se – is the functional driver of the bias. In the over-cancelled condition (purple), the pattern reverses, with biases in the opposite direction. To maintain the full factorial design (Eccentricity **×** Flow Manipulation **×** Heading), we included trials where gaze was centered and straight ahead. Although no natural curl is generated at this zero-eccentricity position, we purposely introduced curl at rotation speeds corresponding to the ‘cancelled’ and ‘over-cancelled’ conditions (simulating random leftward or rightward gaze). As shown in the central panel of Figure 3, this introduced flow induced the predicted biases despite the absence of gaze eccentricity, confirming that the retinal flow pattern, rather than the physical orientation of the eyes, is the primary driver of the perceived heading shift.

**Figure 3:**
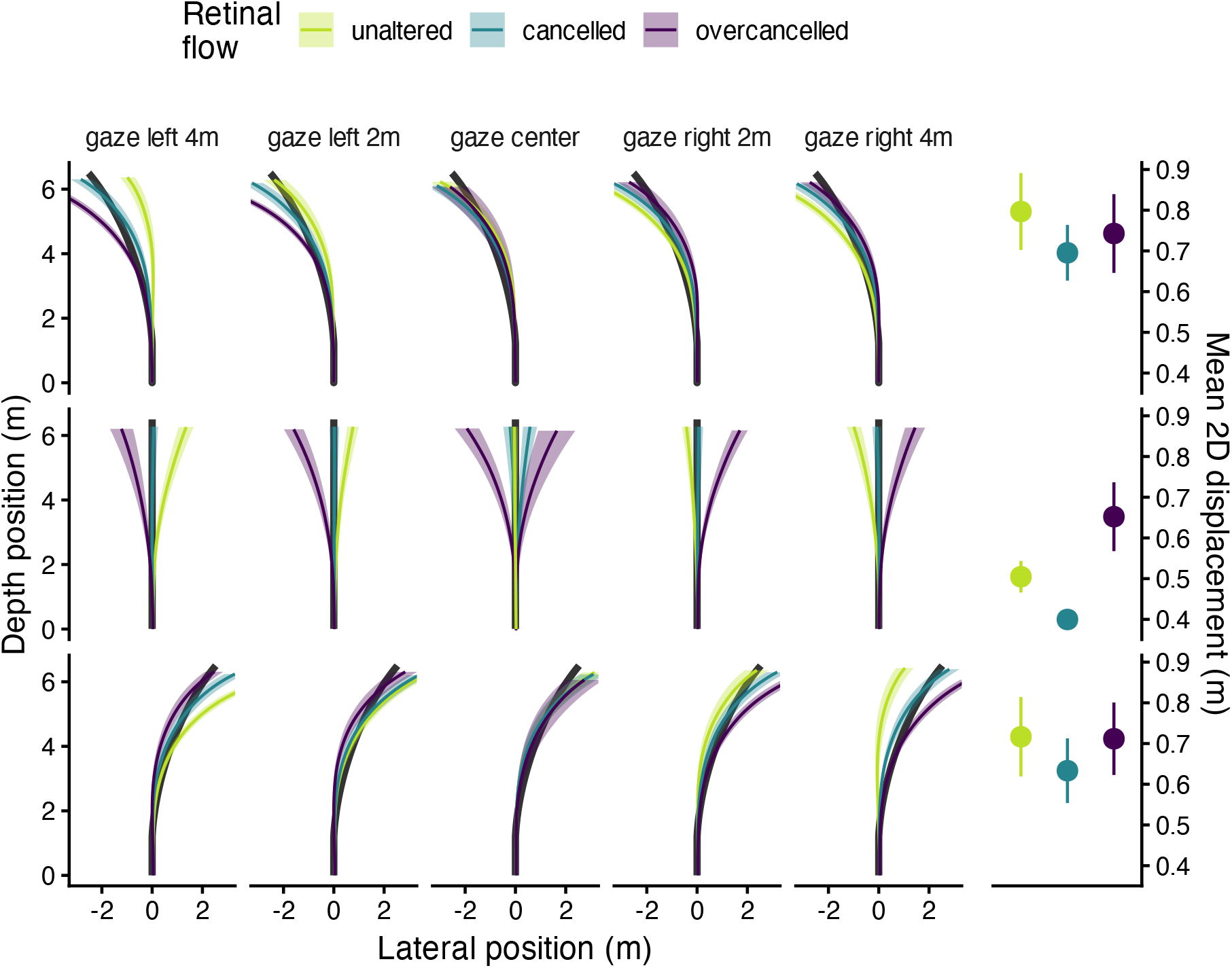
Average perceived heading for each gaze condition (columns) and each physical path curvature (rows), separately for the three retinal–flow manipulation conditions: unaltered curl (green), cancelled curl (cyan), and over-cancelled curl (purple). Shaded envelopes indicate between-observer variability (95%-CI). Note the heading biases in the gaze center condition/middle row. Unexpected positive or negative curl was added in some trials to mantain a full factorial design (see main text for more details). The right axis applies to the last column and illustrates the mean 2D displacement between observed and physical paths. This measurement indicates the mean displacement that is required for the observed path to align with the physical one. The displacement is shown for the different flow manipulations (color-coded). The error bars indicate between-observer variability (95%-CI).

The last column in Figure 3 shows the average 2D displacement (in meters) required for the observed paths to align with the physical ones. This measure gives an idea of the similarity between the reported paths and the physical exposed trajectories in the experiment. In general, the difference between perceived and physical paths is smaller when the path is straight (central panel of last column in Figure 3). Consistent with the reported paths in the different flow manipulation conditions, the displacement is smaller in the cancelled flow curl (color-coded) for all headings (different rows). This resulted in a significant quadratic effect of Flow manipulation, *β* = 0.09, SE = 0.023, t(526) = 3.997, p < .001, indicating that mean deviation was lower in the cancelled condition compared with the unaltered and overcancelled conditions.

### Fitting the controller

To assess how well the curl-based controller accounts for reported perceived headings, we fitted the model as described in the Methods. The fit incorporates the mean image curl computed via Equation 1 from the experimental videos, assuming perfect gaze stabilization on the fixation point. Fits were performed for both the aggregate data and individual participants (see Figure 4 and its corresponding supplemental figures in the SI). As shown in Figure 4, the model with separate parameters per condition captures the average perceived trajectories (thin lines) with remarkable accuracy (thick red lines). In these separate fits, the mean 2D deviation per step was minimal, ranging from 0.066 m in the cancelled flow condition to 0.080 m in the unaltered condition (see Table 1 in Appendix 3). Statistical analysis via ANOVA on the individual fits confirmed that while separate fits yielded a significantly smaller loss than the join approach (*F* (1, 55) = 206.27, *p* < 0.001), flow manipulation itself had only a marginal effect on fit quality (*F* (2, 55) = 2.69, *p* = 0.077). Interestingly, the cancelled condition showed a slightly smaller loss than the other two (Quadratic contrast *t*(55) = 2.356, *p* = 0.022).

**Figure 4:**
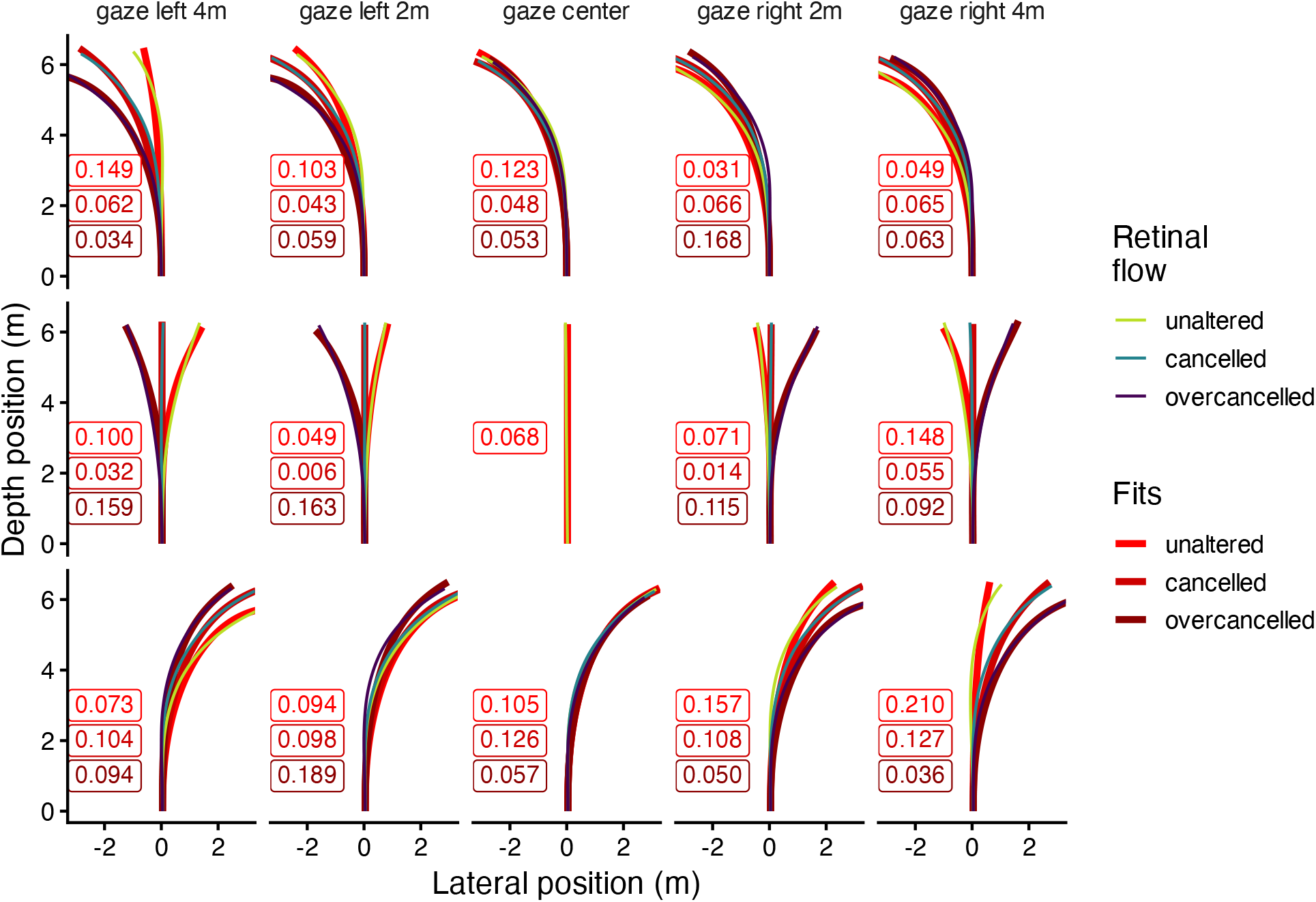
Separate fits of the controller. Average perceived heading across participants for each gaze condition (columns) and each physical path curvature (rows), separately for the three retinal–flow manipulation conditions (thiner solid lines): unaltered curl (green), cancelled curl (cyan), and over-cancelled curl (purple). The different red thicker solid lines denote the best fit of the controller. The numbers in each panel indicate the average lateral deviation per step between the fit and the observed heading. Note that for the centered gaze and straight path, only the unaltered flow condition was fitted.

We also evaluated a joint fit—using only two parameters to account for all heading and eccentricity conditions. We view this joint fit not as a perfect model, but as a deliberate limit test of the curl-based model. In reality, the control “Gain” (the weighting of the curl signal) is likely tuned dynamically based on task demands and sensory confidence. By forcing a single, fixed linear relationship between retinal curl and steering effort, we intentionally ignore this contextual variability to determine how much behavior can be explained by a minimalist controller. Although this compromise naturally results in a larger 2D deviation, it successfully captures the core qualitative trends: a) tracking near-straight trajectories when curl is cancelled; b) reproducing the systematic underestimation of curvature when gaze aligns with the curve; and c) predicting the reversal of bias directions in the over-cancelled condition (purple lines). Upon closer analysis, the most pronounced discrepancies in the joint model occur in this ‘over-cancelled’ condition, where sensory flow evidence becomes ecologically inconsistent with extra-retinal gaze direction, likely requiring complex sensory re-weighting. We therefore frame this joint model, not as a perfect model, but as a parsimonious, first-order approximation. Despite the higher deviation inherent in fixing parameters, its statistically superior (lower) AIC measures (Table 1 in Appendix 3) demonstrate that a minimal, unified curl-based mechanism provides a fairly good approximation for heading perception across diverse conditions.

### Neural simulations

Having established that a curl-based controller successfully accounts for heading responses, we next sought to determine how this mechanism might be implemented in a neurophysiologically plausible way. To do so, we developed a recurrent ring network model designed to transform local foveal flow into global heading estimates (see appendix 2). This architecture explicitly incorporates the role of active gaze by modeling the foveal curl as a localized inhibitory drive on the network’s activity. Because our primary aim with this neural model is to mechanistically explain the origin of the systematic opposite-gaze bias, we focused our simulations exclusively on the straight-path conditions. The model processed optic flow directly from the experimental video sequences—assuming perfect gaze stabilization—and simulated the continuous neural dynamics from local motion extraction to the linear decoding of the population vector.

To identify the specific drivers of the heading bias, we systematically explored the network’s parameter space. We varied the prior strength (*I*_0_ ∈ [0.03, 0.24]), the prior width (*σ*_*p*_ ∈ [0.18, 0.25]), the spatial extent of gaze inhibition (*σ*_*k*_ ∈ [0.02, 0.04, 0.08, 0.12]) and the inhibitory gain (*K*_*p*_ ∈ [0.4, 0.8]). Table 1 in the appendix 2 shows the range for the different parameters used in the simulations.

#### Simulation results

The network dynamics provide a clear mechanistic explanation for the systematic heading bias observed during eccentric gaze. In Figure 5, we observe the network components for a trial in which the gaze direction is 4 m to the left during forward translation. Figure 5A shows the recurrent connectivity following a standard Mexican hat profile, characterized by local excitation and broader surround inhibition. The activity bump does not settle at the objective straight-ahead (0^°^) but stabilizes at a positive (rightward) heading offset, as illustrated in Figure 5B. This displacement is driven by the competition shown in the mechanism panel (Figure 5C): the gaze-centered inhibitory drive (red dashed line) suppresses the leftward portion of the ring, effectively ‘pushing’ the neural activity bump (black line) away from the gaze location and to the right of the straight-ahead prior (blue dotted line). The Phase Portrait (Figure 5 D) confirms this as a stable state; the drift dynamics 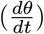 converge toward a stable fixed point where the curve crosses zero at a rightward offset. Together, these results demonstrate how localized inhibition, within a parietal ring attractor, transforms gaze-contingent sensory signals into a shifted global heading estimate. For results across a range of eccentricities during forward translation, see Figure 5 Suppl. Figure 1 in the SI. The presence of temporal oscillations in the activity bump when gaze is aligned with the direction of travel (Figure 5 Suppl. Figure 1 B) or at low eccentricities (A and C) represents a state of high competition within the attractor. As seen in Figure 5 Suppl. Figure 1 B, when the gaze-centered inhibition —the sensory *pull*— overlaps directly with the straight-ahead prior —the internal *push*— it creates a central suppression that counteracts the recurrent excitation of the network. This conflict results in a concave activity profile and rhythmic fluctuations in bump intensity as the network continuously attempts to re-stabilize the representation against the central inhibitory zone. Notably, these oscillations persist when gaze is less eccentric (e.g., at -2 and 2 m in rows A and C respectively), as the proximity of the inhibitory signal to the prior creates a similar destabilizing interaction. Despite these internal dynamics, the decoded heading remains accurate due to the symmetry of the competing forces, suggesting that while representation coherence may decrease, the population vector readout remains robust to central sensory-prior conflict.

**Figure 5:**
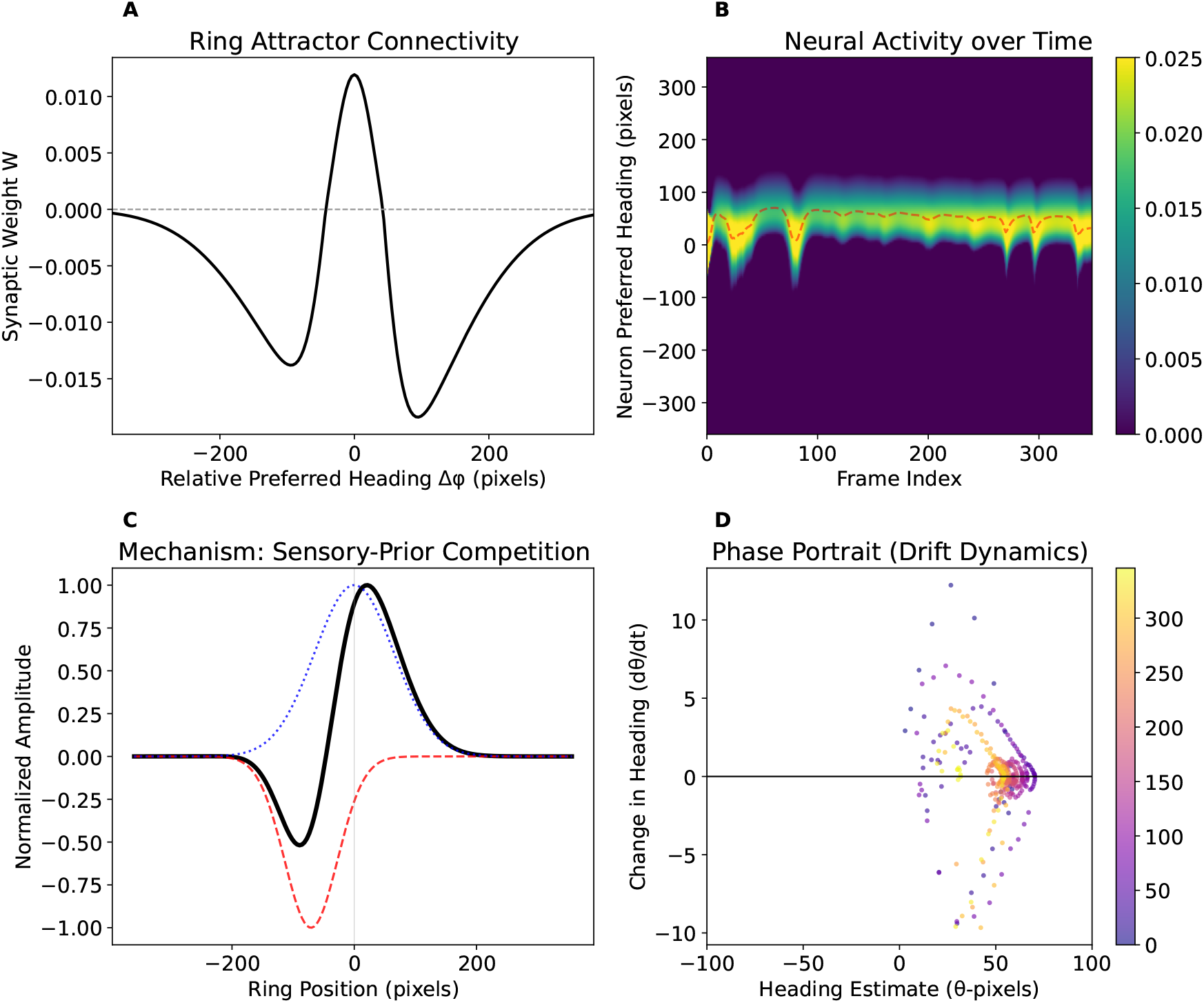
Neural network model of gaze-contingent heading bias. The parameters used to produce these panels are: *I*_0_ = 0.03, *K*_*p*_ = 0.4, *σ*_*p*_ = 0.18, *σ*_*k*_ = 0.12. (A) Ring attractor connectivity showing synaptic weight (*W*) as a function of relative preferred heading (Δ*ϕ*) in pixels, featuring a central excitatory peak and asymmetric inhibitory sourround. While we present results using asymmetric connectivity, no significant differences were observed between asymmetric and symmetric configurations. (B) Heatmap of neural activity across neuron preferred headings (y-axis) over the frame index (x-axis), with a dashed red line indicating the decoded population heading estimate for the trial in which gaze directed 4 m to the left. (C) Mechanism of sensory-prior competition plotting normalized amplitude against ring position in pixels, illustrating the spatial alignment of the straight-ahead prior (blue dotted line), gaze-centered inhibition (red dashed line), and the resulting neural activity bump (black solid line). (D) Phase portrait showing the change in heading (*dθ*/*dt*) versus the heading estimate (*θ*) in pixels, with a horizontal line at zero marking the convergence of the trajectory toward a stable fixed point The color bar denotes frame number.

From the neural model heading estimates (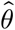, in pixels) we reconstructed the 3D paths using a pinhole camera model (see Heading Estimation and 3D Path Reconstruction section in SI) that are shown in Figure S17 of the SI. This figure demonstrates that the observed instant headings pattern shown in Figure 2 is best captured by a network configuration featuring a weak straight-ahead prior (*I*_0_ ≈ 0.03) and moderate inhibitory gains (*K*_*p*_ between 0.4 and 0.8). Within this parameter space, the localized, gaze-contingent inhibition effectively shifts the population activity bump, reproducing the characteristic lateral spread seen in the behavioral data. A low prior value (*I*_0_ = 0.03 in the simulation grid) is essential. Higher prior strengths (e.g., *I*_0_ = 0.24) overly anchor the heading estimate to the center (0 m), failing to produce the significant lateral spread observed in the human data. *K*_*p*_ = 0.4 closely matches the tighter trajectory fan seen in the top central panel of the empirical data, while *K*_*p*_ = 0.8 captures the wider lateral deviations observed in the bottom central panel, where the bias is more pronounced. Concerning the inhibition width (*σ*_*k*_), values between 0.08 and 0.12 provide the most realistic structural match. Very low values (*σ*_*k*_ = 0.02, 0.04) produce negligible bias even at high gain, as the localized inhibition does not sufficiently overlap with the heading activity bump to drive a shift. The prior width value of *σ*_*p*_ = 0.18 appears to provide a more sensitive range for capturing the gaze-contingent shift compared to the wider *σ*_*p*_ = 0.25. These results suggest that the interplay between sensory-driven inhibition and a stabilizing straight-ahead prior within a standard Mexican-hat recurrent architecture is sufficient to account for the gaze-contingent biases observed in human heading perception.

## Discussion

### Heading Bias Reveals Retinal Curl as a Control Variable

We provide evidence that mean retinal curl plays a functional role in heading perception, as manipulating retinal flow predictably alters perceived trajectories. This finding suggests that curl could be used as a control signal rather than being a “nuisance” component of the optic flow that must be filtered out. While previous work has emphasized the decomposition of flow into translational and rotational components to recover heading (Beintema et al., 2004; Heeger and Jepson, 1992; Longuet-Higgins and Prazdny, 1980), we show that the visual system exploits the curl generated by gaze stabilization to perceive heading, as the sustained presence of this image curl predictably biases heading judgments. When the naturally occurring curl was counteracted, the systematic heading bias disappeared. This confirms that the bias is not an artifact of gaze eccentricity itself, but a direct consequence of the underlying flow geometry due to sustained gaze stabilization. In addition, the flow manipulation reveals a direct link between the amount of curl and perceived heading: not only did the bias opposite to fixation disappear when the curl was cancelled but also the bias re-appeared towards fixation when the curl was over-cancelled (see paths in the middle row in Figure 3). This manipulation led to counter-intuitive findings like the reported path was closer to the physical/experimental one in the cancelled condition (last column in Figure 3)

This is consistent with Matthis et al. (2022), who observed that the head-centered FoE during natural walking is too variable to guide navigation reliably. Instead, they found that the magnitude of retinal curl provides a stable, gaze-relative trajectory signal. By experimentally isolating this curl, we provide the perceptual link demonstrating its direct effect on perceived heading. Thus, curl acts as a functional surrogate; when combined with extra-retinal gaze signals, it allows the visual system to bypass the unstable FoE in favor of a robust, rotation-based control signal.

### Reference frames

Heading perception has traditionally been studied using either retino-centric (Grigo and Lappe, 1999; Rieger and Toet, 1985; Stone and Perrone, 1997) or world-centered (Banks et al., 1996; Royden et al., 1992; van den Berg, 1992; van den Berg and Brenner, 1994; Warren and Hannon, 1988) reference frames. Although reporting heading in world coordinates often feels more intuitive to observers, this extrinsic frame likely reflects the specific demands of the experimental task rather than the coordinate system of the underlying sensory information. Retinal curl inherently provides a gaze-centered signal, specifying the angular offset between the observer’s trajectory and their gaze axis. To fulfill the task requirement of a world-centered judgment, observers could simply anchor the known gaze direction to the 3D scene (e.g., relative to the initial longitudinal alignment, *θ*_0_ in Figure 1 B) and apply this curl-derived offset. Thus, observers can resolve an extrinsic trajectory without ever needing to extract a world-fixed FoE.

While this gaze-relative offset explains our data, we must consider whether an alternative, egocentric reference frame could also account for the observed opposite-to-gaze bias. Typically, gaze-dependent heading judgments bias *toward* the direction of gaze (Grigo and Lappe, 1999; Royden et al., 1992). To explain an *opposite* bias using an egocentric frame or visual direction (Rushton et al., 1998; Wann and Land, 2000), one must assume that retinal curl induces an illusory body rotation. For instance, if rightward fixation is misinterpreted as rightward body rotation, the perceived straight-ahead drifts right, causing a straight physical path to be erroneously judged to the left. However, this illusory shift is highly unlikely to explain our results for two reasons. First, egocentric visual direction relies on matching discrete targets or end-points (Rushton et al., 1998), whereas our continuous, landmark-free ground plane strongly favored flow-based processing (Wilkie and Wann, 2003). Second, interpreting foveal curl as a global body rotation creates a fundamental conflict between the exposed optic flow and the expected visual consequences of actual body rotation.

A second alternative explanation for the opposite-gaze bias lies in the under-compensation of extra-retinal signals (Freeman et al., 2010). If the brain systematically underestimates the magnitude of an eccentric eye movement, the perceived gaze is located closer to the mid-line than its true physical position. Adding an accurate curl-based visual offset to this under-compensated gaze vector would push the resulting perceived heading past the actual trajectory, generating an opposite-gaze bias. However, this extra-retinal under-compensation cannot explain the bias in our central gaze condition, where participants maintained fixation directly straight ahead while structured foveal curl was artificially inoculated into the display.

### Temporal integration scales

The perception of a curved path during physically straight motion is not new (Grigo and Lappe, 1999; Royden et al., 1994, 1992; Royden, 1994; van den Berg and Brenner, 1994). Typically, longer presentation times facilitate more accurate heading judgments (Longuet-Higgins and Prazdny, 1980; Royden, 1994; Stone and Perrone, 1997; Xie and Li, 2025). However, despite our substantial trial durations (>10 s), we observed a robust bias that cannot be attributed to a lack of integration time. Notably, while Grigo and Lappe (1999) found that increasing stimulus duration (from <1 s to 3 s) increased biases toward the direction of gaze, our findings reveal a bias in the opposite direction. This difference suggests that the nature of the bias in our study and those previously reported must obey different underlying mechanisms.

A key difference is the rotation threshold. Earlier biases emerged only at simulated rotations exceeding 1°/s, with perception remaining accurate at the lower velocities used here (Warren and Hannon, 1990). Additionally, because previous studies used brief 2–3 second stimuli, they missed that the opposite-gaze bias requires 3–5 seconds to fully manifest (Figure 2). This delay aligns with optic flow stabilization times (Warren et al., 2001) and suggests the brain integrates retinal curl over several gait cycles. Such extended processing matches evidence that global motion signals (e.g., expansion and rotation) require 1–3 second integration windows (Burr and Santoro, 2001), far surpassing the ∼200 ms limit of local motion.

While instantaneous flow is critical for millisecond-scale postural balance (Bardy et al., 1999; Powell et al., 2026) and “instantaneous” retino-centric heading judgments asymptote around 400 ms (Perrone and Stone, 1994; Stone and Perrone, 1997), locomotor path control relies on a much longer integration scale. Such brief windows can be insufficient to resolve the ambiguities inherent in curved paths. Indeed, recent evidence confirms that longer stimulus durations improve heading judgments by stabilizing the trajectory estimate, rather than by enhancing the flow-parsing process itself (Xie and Li, 2025). Our use of continuous reporting—unlike the binary, post-trial responses of most prior studies—allowed us to reveal a temporal evolution like in steering tasks (e.g., Warren et al., 2001). These continuous dynamics suggest the visual system relies on time-varying optic flow (Burlingham and Heeger, 2020) to perceive a “future path” (Cutting et al., 1992; Li and Cheng, 2011), rather than extracting a momentary heading vector from instantaneous flow (Perrone and Stone, 1994; Stone and Perrone, 1997). Through sustained gaze stabilization, the brain moves beyond immediate transients, integrating foveal signals into a stable representation of the upcoming trajectory. Readers can experience this temporal build-up firsthand: maintaining a sustained lateral gaze while walking in an open space gradually induces a subtle, integrated drift that eventually becomes a clear departure from the intended path.

### Active Steering vs. Heading Recovery

Navigation research distinguishes passive heading perception from active steering control (Goodridge et al., 2023; Powell et al., 2024; Wilkie and Wann, 2003). Traditional models assume explicit heading recovery (e.g., locating an instantaneous FoE) is required for steering, but our controller demonstrates that a trajectory can be maintained simply by nulling the error signal derived from retinal curl. Kim and Turvey (1999) and Wann and Swapp (2000) proposed a conceptually similar strategy that relies exclusively on sensitivity to retinal flow curvature in order to linearize the flow field. However, Saunders and Ma (2011) reported evidence against this pure linearization approach. Unlike this approach, our model does not depend on flow acceleration or curvature alone. Instead, it temporally integrates mean retinal curl, which is disambiguated by extra-retinal gaze variables. As derived in Appendix 1, raw curl becomes a precisely calibrated steering command only when scaled by gaze orientation, eye height, and translation speed (Frenz and Lappe, 2005). This gaze-scaled curl allows the visual system to continuously regulate the geometry of a stable “future path” without extracting a momentary heading vector (Tuhkanen et al., 2021; Wann and Swapp, 2000; Wilkie and Wann, 2006).

To validate this mechanism, we applied our controller to the paradigms of Wilkie and Wann (2003) (Experiment 3), successfully matching their reported empirical steering paths (Appendix 3). While they proposed a model integrating retinal flow with egocentric cues like visual direction (Rushton et al., 1998), our empirical findings demonstrate that flow dynamics can completely override egocentric signals. Even when an eccentric fixation point provided a strong, constant visual direction cue, cancelling the retinal curl eliminated the steering bias entirely, while our “over-cancelling” condition reversed its direction. This reversal contradicts a primary reliance on the visual direction of the target. Instead, it suggests that in the absence of structural landmarks—and in the presence of reliable, large-field (>90°) optic flow—the visual system strongly favors gaze-scaled flow processing over visual direction (Wilkie and Wann, 2003).

### Re-evaluating the Focus of Expansion (FoE)

The concept of the Focus of Expansion (FoE) was originally proposed by Gibson in the context of aircraft landings—a scenario characterized by minimal eye or head rotation (Gibson, 1950). However, the relevance of a translational head-centred FoE in natural contexts has been recently questioned (Matthis et al., 2022; Muller et al., 2023). While our results support the view that explicitly extracting a pure, de-rotated FoE is not required for steering, this does not render the FoE useless. The FoE likely remains highly relevant during high-speed locomotion, where gaze shifts are less frequent relative to the speed of travel, leading to a less disrupted flow field (Muller et al., 2023). Furthermore, even in complex scenes characterized by relative object motion, *pseudo-FoE* signals have been shown to drive heading biases (Layton and Fajen, 2016). We speculate that rather than subtracting global rotation to uncover a hidden FoE, the visual system exploits these uncorrected pseudo-FoE locations directly, combining this shifting retinal center of motion with extra-retinal signals and foveal curl to resolve the trajectory.

This perspective aligns closely with the neural model of MSTd proposed by Layton and Browning (2014), while our ring-attractor model extends this framework into a downstream parietal decoding mechanism. Layton and Browning (2014) suggested that MSTd integrates rotation directly rather than treating it as a nuisance component. In their model, MSTd hypercolumns represent self-motion simultaneously through the spirality (curl) of the most active units and the visuotopic location of this activity peak (the pseudo-FoE). Building upon this MSTd-like extraction of spiral motion (Duffy and Wurtz, 1991; Graziano et al., 1994), our neural model provides a biologically plausible bridge to motor control. We propose that downstream parietal regions, such as VIP (Schaafsma and Duysens, 1996) and 7a (Read and Siegel, 1997), to which MSTd projects (Born and Bradley, 2005), utilize a ring-attractor network to interpret this sensory output. In our model, standard center–surround (Mexican-hat) recurrent connectivity interacts with a weak straight-ahead prior and gaze-centered inhibition. These inherent push–pull dynamics effectively transform localized sensory inhibition into a global shift of the activity bump, naturally generating the psychophysical biases observed in our data. Furthermore, the network’s recurrent dynamics act as a low-pass filter, smoothing high-frequency, gait-related temporal oscillations into stable trajectory estimates. Together, these frameworks demonstrate how standard cortical architectures can seamlessly transform uncorrected retinal flow into robust steering commands, without ever explicitly extracting a pure FoE.

### Generalizability and Testable Predictions

Our results align with a growing consensus in sensorimotor research that incidental sensory signals often carry vital functional information (Rolfs and Schweitzer, 2022). Much like saccade-induced motion streaks facilitate gaze correction (Schweitzer and Rolfs, 2021), or oculomotor cycles actively format spatiotemporal visual input (Boi et al., 2017), retinal curl appears to be a systematic byproduct of gaze stabilization that the visual system exploits for navigation. Under this view, the biases observed in 3D environments are not failures of calculation, but rather the signature of a proportional controller shifting perceived heading toward a “null-curl” orientation.

This control-law perspective leads to several testable predictions. First, in environments with low visual texture or visibility (e.g., fog), the reliability of the curl signal should decrease, forcing the system to rely more heavily on “straight-ahead” internal priors and thus reducing the magnitude of the induced bias. Second, because our model relies on the precise calibration between extra-retinal signals and retinal curl, any perturbation of eye-position signals—whether through experimental manipulation or clinical conditions—should result in systematic steering errors that match the mis-scaling of the curl signal. Finally, while our focus was on ground-plane fixation, the model predicts that any sustained gaze strategy that introduces structured rotation—such as tracking a moving agent or a point on a vertical wall—should generate predictable trajectory biases dictated by the specific geometry of the resulting foveal flow.

## Conclusion

We conclude that the rotational component of optic flow (curl), generated during gaze stabilization, is an actively used signal to control heading. It acts as a navigational cue rather than noise, as evidenced by the elimination of steering biases when curl is experimentally cancelled. Our findings challenge the necessity of explicitly extracting the Focus of Expansion for online control of locomotion. The observed behaviors are supported by a neural model based on established properties of motion processing areas. The interaction between sensory flow inputs and internal priors within a recurrent network suffices to explain the gaze-contingent biases observed in our experimental data.

## Methods

### Participants

We tested 12 participants (five self-identified men and seven self-identified women), aged between 24 and 59 years (mean: 30, SD: 9), all with normal or corrected-to-normal vision. Except for one, all participants were naïve to the aims of the study and volunteered to take part. The study forms part of an ongoing research program approved by the Ethics Committee of the University of Barcelona (IRB 00003099) and conducted in accordance with the principles of the Declaration of Helsinki.

### Displays and conditions

Participant motion was simulated as a translation parallel to the ground plane at a sustained walking speed of approximately 1 m/s. This motion incorporated characteristic bounce and swing components derived from a single gait profile (see Fig. S1 in the SI). This profile was recorded once by an independent individual wearing an HMD tracker while walking along the predefined experimental paths in a virtual environment, and then was applied to all participants to ensure stimulus consistency. While we acknowledge that using a standardized gait profile may introduce individual-level variability in perception, as the simulated motion may not align perfectly with each participant’s unique gait dynamics, this approach was chosen to ensure that the reported perceptual biases are a robust consequence of the experimental variables (gaze direction and retinal curl) rather than artifacts of varying motion kinematics.

Each trial lasted between 11 and 12 s.

The ground plane consisted of a 50 × 50 m surface mapped with a naturalistic texture (see Fig. 1A) generated from simplex noise patterns whose spatial power spectrum followed a 1/*f* ^2^ distribution. The temporal frequency of these patterns generated by simulated self-motion is consistent with the statistics of natural videos as described by (Dong and Atick, 1995) also following a 1/f-type temporal power spectra (exponent of 2). The textures were created in real time using OpenGL shaders within a custom Python program, designed for computational efficiency and allowing online manipulation of the texture in specific experimental conditions (see *Flow manipulation conditions* below).

The experiment was run on an Intel i7-based workstation (i7-9700F, Intel, Santa Clara, CA, USA) equipped with an NVIDIA GeForce RTX 2060 SUPER GPU. Images were rendered at 120 Hz with a resolution of 1920 × 1080 pixels and displayed monocularly via a PROPixx projector (VPIxx Technologies, Saint-Bruno, QC, Canada) onto a back-projection screen (2.03 m × 1.16 m), viewed from a distance of 1.0 m, resulting in a visual field of approximately 91º. The scale of the visual stimulus was precisely matched to the simulated environment by aligning the virtual camera’s field of view with the physical geometry of the screen, ensuring a veridical 1:1 mapping between virtual and physical space.

To verify fixation, eye movements were recorded with a Pupil Labs Core (Berlin, Germany) eye tracker operating at 200 Hz. Trials were discarded if the median distance between gaze position and the fixation point exceeded 3º.

#### Path conditions

Participant trajectories could be either straight (length of about 6.5 m) or curved to the left or right (see Fig. 1B). The curved paths resulted in a final heading of ±45° relative to the initial heading (0°) in world coordinates. The three path types (straight, left, and right) were interleaved on a trial-by-trial basis.

#### Eccentricity conditions

In all trials, a fixation point (yellow dot in Figure 1A) was presented on the ground and remained fixed in world coordinates. Relative to the participant’s initial position (x = 0, z = 0) and heading (0°), the fixation point could appear at one of five lateral positions or eccentricities, x = {-4, -2, 0, 2, 4} m, and was initially located 20 m ahead (see colored dots in Fig. 1B). As simulated self-motion progressed, the fixation point appeared to approach the observer, necessitating a gradual increase in gaze angle. Participants were not explicitly instructed to use specific eye or head movements to maintain fixation; instead, they were permitted to engage the head-eye system naturally to track the target. Under perfect fixation, the expected head/eye rotation rate increased for the largest eccentricity from about 0.4 º/s to 0.8°/s (see Fig. S2 in the SI). The eccentricity condition was randomized across trials.

#### Flow (curl) manipulation conditions

When fixating a stationary eccentric point while translating straight ahead, the retinal image contains an expected rotational component— referred to as curl—around the fovea. This effect is illustrated by the flow lines in Fig. 1A, which shows a snapshot of the retinal image consistent with walking straight ahead while fixating a point located to the left (positive curl). Figures 1C–E display the mean curl distributions for fixation points at varying eccentricities: leftward positions (−4 m, −2 m) produced positive curl, the center (0 m) resulted in near-zero curl, and rightward positions (2 m, 4 m) produced negative curl.

To compute the curl shown in Figs. 1C–E, we simulated experimental trials (videos downsampled to 800 × 452 px at 30 Hz) and computed optic flow using the Farnebäck algorithm (OpenCV). To ensure robust measurements and alleviate texture-dependent variability that can be introduced by the Farnebäck algorithm, we ran each trial 10 times using different textures always consistent with natural image statistics. For each frame *t*, we extracted the 2D image curl from the horizontal (*f*_*x*_) and vertical (*f*_*y*_) flow components:

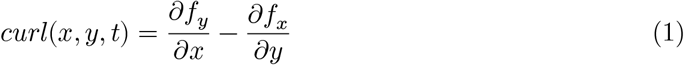

Beyond this algorithmic extraction, the retinal curl magnitude is geometrically determined by the observer’s translational speed (*v*), eye height (*h*), and gaze orientation (pitch *α* and yaw *ψ*). As derived in Appendix 2, for a ground plane this relationship approximates to:

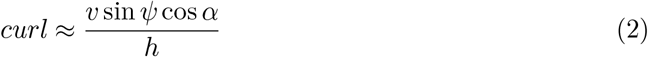

We computed the mean image curl 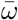 for each frame. Since the visual scene included gait-related oscillations (affecting the curl through both pitch and yaw), the raw computed curl was temporally smoothed (temporal window of 2.4 s). This processed signal was then used to make predictions based on the controller, ensuring robustness to high-frequency head movements. These values approximate the mean retinal curl expected under accurate fixation and define the *unaltered curl condition*.

In some conditions, this expected curl was counteracted (*cancelled curl condition*) by adding an equal and opposite rotational component centered on the fixation point. Figure 1C,E also show the corresponding curl histograms (lighter colors), which are centered at zero—matching the distribution obtained when fixating straight ahead in the direction of motion (mean zero curl, Figure 1D). In an additional set of trials, the imposed counter-rotation was doubled to produce an *overcancelled condition*, in which the curl was reversed beyond neutralization (histograms not shown in Figure 1).

### Procedure

Participants continuously reported their perceived self-motion direction in the 3D scene while maintaining fixation on a target dot. Responses were collected via a custom rotative encoder (0.3° angular resolution) configured as an intuitive steering wheel interface, which participants reported was very easy to operate. To ensure high-frequency, non-blocking data acquisition, the device was interfaced through an Arduino Uno. We utilized a dedicated Python thread for direct serial communication, allowing for real-time polling of the encoder state without interfering with the primary stimulus rendering loop.

At the beginning of each session, participants completed a five-point calibration procedure for the eye tracker. Each session comprised 45 trials (3 trajectory types × 5 fixation positions × 3 flow manipulation conditions), and each participant completed a total of 10 sessions.

### Computation of perceived path

To reconstruct the participant’s estimated path from the angular response, we treated the reported heading angle *θ*_*t*_ (radians) as specifying the instantaneous lateral slope of the perceived trajectory at frame *t*. For each time step, we computed the incremental simulated forward displacement

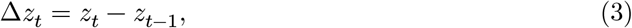

and converted the response angle into a lateral increment via

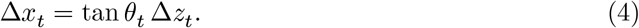

The estimated lateral position was then obtained by forward integration starting from the initial position *x*_0_:

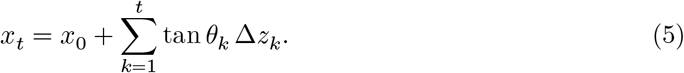

This yields the sample-wise reconstructed lateral trajectory *x*_*t*_ consistent with the participant’s angular responses.

### Control Model

We interpret the reported path as the output of a curl–driven feedback mechanism inspired by point attractor models used previously in steering control (Fajen and Warren, 2007; Wilkie and Wann, 2003). Let *θ*_*t*_ denote the heading direction (direction of forward velocity in world coordinates) and *ψ*_*t*_ the gaze direction. We consider their difference

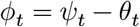

as a instantaneous heading error. The mean optic–flow curl 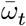 is taken as a sensory measurement of this discrepancy, so that 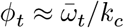, with *k*_*c*_ being a curl–to–angle scale factor. This is motivated by the fact that curl increases whenever there is a discrepancy between *θ* and *ψ*. The observer controls heading by applying a yaw rate input 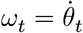:

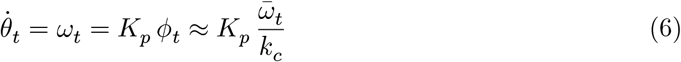

so that positive curl induces a leftward turn (increasing *θ*) and negative curl induces a rightward turn. If gaze is held fixed (e.g. steady fixation), 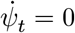 and therefore

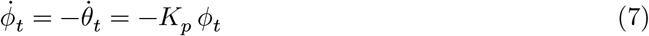

This is a simple first-order linear differential equation, and the heading error decays exponentially with a correction time constant 1/*K*_*p*_:

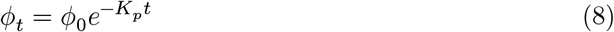

Thus, heading is continuously steered toward gaze until curl vanishes. In our reconstruction we integrate this controlled heading forward in time to generate the perceived trajectory implied by the observed curl (see video S3 in the SI).

### Trajectory fits

To test if perceived trajectories can be accounted for by the curl signal, we used the controller-defined in Equation 6 which we integrated forward with an explicit Euler step to obtain the estimated heading *θ*. With sampling interval Δ*t*, constant forward speed (*v*), and initial state (*θ*_0_, *x*_0_, *z*_0_) taken from the first sample of each trial, the discrete update is

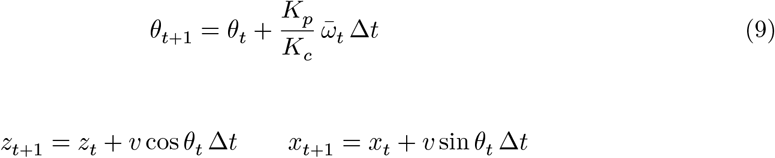

This generates a predicted 2D path 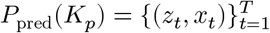 from the observed mean curl 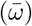 sequence. We then estimate *K*_*p*_ and forward velocity (*v*) as free parameters by minimizing a time-warped path discrepancy (*J*) using Dynamic Time Warping (DTW)(Giorgino, 2009) on the (*z, x*) sequences between the predicted and perceived paths:

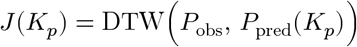

where

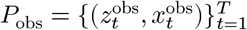

is the observed path.

We fix *K*_*c*_ (Equation 6) to 1, absorbing the unknown curl-to-angle scale into *K*_*p*_.

We used two fitting approaches: (1) **independent** or **separate** fits, in which parameters were fitted independently for each combination of heading, gaze eccentricity, and retinal flow manipulation and (2) **join fits**, in which a single set of parameters is used across all heading and gaze eccentricity conditions but they were different for each flow manipulation. Each approach was tested with two parameters (*K*_*p*_, and the forward velocity, *v*). The motivation for including forward velocity in the fit was to compensate for variations in the timing of perceived heading responses. For the separate fits, the number of parameters was 30 (for the 2-parameter approach, *K*_*p*_ and *v*). In contrast, the joined approach used only 2 parameters for all heading and eccentricity conditions.

In order to compare the different models, we employed information criteria adapted for distance-based model comparison. For each model, we computed a surrogate log-likelihood based on the normalized alignment cost:

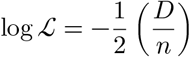

where *D* represents the total DTW distance and *n* the number of observations, with *D*/*n* representing the average alignment cost per observation. This normalization ensures appropriate scaling for information criteria computation. We then computed the Akaike Information Criterion as:

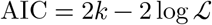

where *k* represents the number of parameters.

## Data availability

The data supporting the findings of this study and code (R and python) are available in OSF at [osf link](https://osf.io/b37rg/overview?view_only=35b748a234b7459a9e9df1b41a2d4a44).

## Acknowledgements

We thanks Cristina de la Malla for her helpful comments and suggestions on the manuscript.

## Appendix 1

### Derivation of curl

Let’s set up an eye-centered coordinate frame (*X, Y, Z*), where the eye is at the origin and the *Z*-axis is the line of sight (gaze direction) pointing directly at the fixation point.

- The observer moves with forward velocity *v* along the world heading.
- The fixation point is on the ground at 3D distance *d* from the eye and eye height is *h*.
- The gaze is offset from the heading by a yaw angle *ψ* horizontally and pitched downward by an angle *α*, where sin 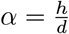.
- Projecting the forward velocity *v* into our eye-centered frame gives the eye’s translational velocity (*v*_*x*_, *v*_*y*_, *v*_*z*_):
- *v*_*x*_ = −*v* sin *ψ*
- *v*_*y*_ = −*v* sin *αα* cos *ψ*
- *v*_*z*_ = *v* cos *α* cos *ψ*

To find retinal flow, we need inverse depth 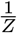 as a function of the image plane coordinates (*x, y*). We assume that the ground plane is perfectly horizontal in the world, but in our eye-centered frame (which is pitched down by *α*), the equation of the ground plane becomes: *Z* sin *α* − *Y* cos *α* = *h*

Dividing by *Z* and substituting the image coordinate 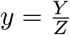, we get the inverse depth profile of the ground:

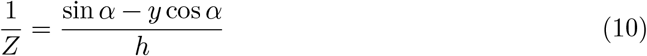

Notice that looking vertically in the image plane changes the depth. The gradient of inverse depth at the fovea (*x* = 0, *y* = 0) is:

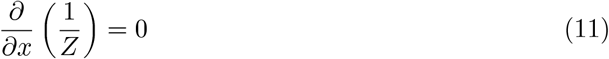

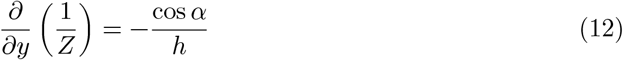

#### Calculating the Curl of the translational flow

The standard optic flow equations for translation on the image plane (*f*_*x*_, *
f*_*y*_) are:

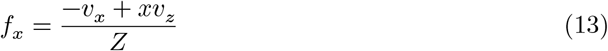

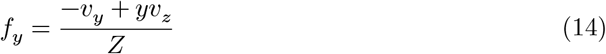

The 2D curl at the fovea is defined as

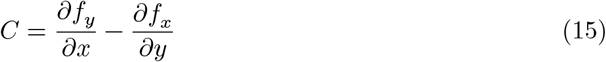

Evaluating this at the origin (*x* = 0, *y* = 0):

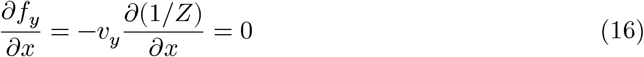

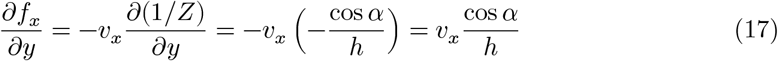

Subtracting the two gives the translational curl:

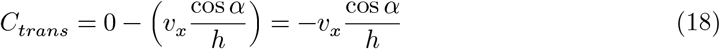

Substituting our earlier *v*_*x*_ = −*v* sin *ψ* into equation Equation 18 we obtain the final expression for curl:

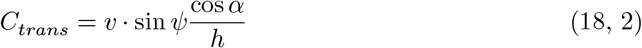

## Appendix 2

### Neural Network Model of Gaze-Contingent Heading Bias

We present a neural model that provides a neurophysiologically plausible implementation of the controller. The model demonstrates that the bias emerges from the interaction between gaze-modulated visual flow processing and a weak straight-ahead prior in parietal heading representation, implemented through standard cortical connectivity. After presenting the model, we show different dynamic aspects of the model and reproduce the bias for the straight ahead trajectories looking at different eccentricities.

#### Local Motion Encoding

Visual motion is encoded by direction-selective units analogous to primate MT cortex, with 8 directional preferences:

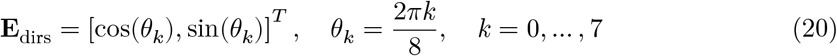

The response of each direction channel at image-plane position **p**_*i*_ = (*x*_*i*_, *y*_*i*_) is given by rectified cosine tuning (Simoncelli and Heeger, 1998):

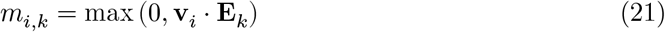

where **v**_*i*_ is the local optical flow vector. In our implementation, optic flow **v**_*i*_ is computed, as before, using the Farnebäck algorithm as a substitute for early visual processing (V1/MT complex). This provides the motion signals that the visual system would extract through V1 to MT processing. The model itself begins with the input to the 8-sector MT-like directional encoding (Equation 21).

#### Curl Computation (MSTd)

To quantify the rotational (curl) component of optic flow around the current gaze position, we project local flow vectors onto the tangential direction of the circle centered at gaze. For each sampled location **p**_*i*_, we first compute its position relative to the gaze point **g**, also encoded in image coordinates (*x*_*i*_, *y*_*i*_):

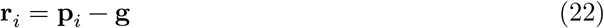

This vector points from the gaze location to the sample point. The direction orthogonal to this vector corresponds to the local tangential direction of rotation around gaze. We obtain this unit tangential vector by rotating **r**_*i*_ by 90° and normalizing:

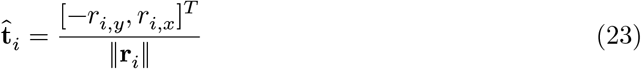

Finally, the contribution of the optic flow at that point to local rotational motion is given by the projection of the flow vector **v**_*i*_ onto this tangential direction:

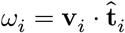

Positive values indicate counterclockwise motion around gaze, and negative values indicate clockwise motion. The mean curl is computed as follows:

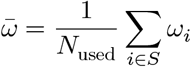

where *S* represents samples satisfying ‖**r**_*i*_‖ > *r*_min_ after trimming outliers. In our implementation, *N*_total_ = 400 samples are drawn within a gaze-centered region.

#### Ring Attractor Dynamics: Parietal Heading Representation

Heading direction *θ* is represented by a ring attractor mechanism consistent with previous studies (Zhang, 1996), with neural activity *x*(*ϕ, t*) at preferred heading *ϕ* evolving as:

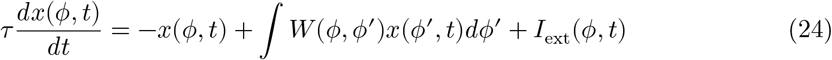

We implement these dynamics by using a discrete ring of *N* = 181 units with preferred headings *ϕ*_*j*_ uniformly spaced between [−*ϕ*_max_, *ϕ*_max_]. The activity dynamics then follow:

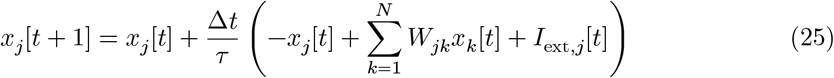

The recurrent connectivity follows a standard Mexican hat profile (Ben-Yishai et al., 1995):

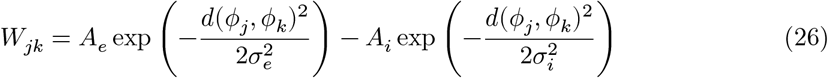

where *d*(*ϕ*_*j*_, *ϕ*_*k*_) is the circular distance between preferred headings. *A*_*e*_ and *A*_*i*_ were set to 1.9 and 1.0 respectively.

#### External Inputs

The external input *I*_ext_(*ϕ, t*) consists of two components:

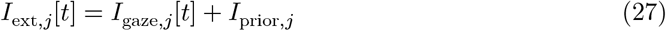

The gaze-modulated inhibitory input is:

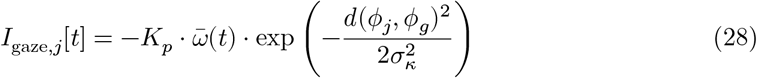

where *ϕ*_*g*_ is the gaze direction in heading coordinates, *K*_*p*_ is the inhibitory gain, and *σ*_*κ*_ controls the width of the gaze-centered modulation, determining how broadly the inhibition spreads around the gaze direction *ϕ*_*g*_

The straight-ahead prior is:

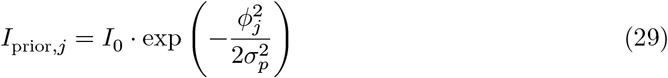

*I*_0_ is the prior strength and *σ*_*p*_ sets the width of the straight-ahead prior, with smaller values producing a sharper preference for *ϕ* = 0

#### Bias generation as Sensory–Prior Competition

Within this framework, the systematic bias opposite to gaze emerges from the interaction between the *gaze-centered inhibitory drive* and a *weak straight-ahead prior* within a stabilizing recurrent Mexican-hat network. A strong prior (*I*_0_ ≳ 0.25) keeps the heading estimate near straight-ahead, while a weak prior *I*_0_ ≈ 0.03 allows sensory evidence to shift the attractor state. The symmetric recurrent dynamics maintain a coherent and stable activity bump. For leftward gaze (*ϕ*_*g*_ < 0) and forward motion 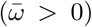: 1) Gaze-centered inhibition creates activity suppression at *ϕ*_*g*_; 2) With weak prior, activity bump shifts toward opposite side and 3) Recurrent dynamics translate inhibition into bump displacement.

The heading direction is decoded using population vector readout:

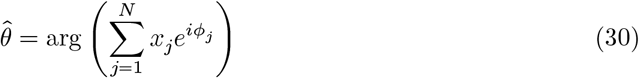

where *N* is the number of ring units (*N* = 181). Unlike some heading perception models (Beintema et al., 2004; Lappe and Rauschecker, 1993), we do not apply sigmoid nonlinearities to neural activities prior to decoding but a linear readout. Gaze-contingent bias emerges even with linear decoding, suggesting it is a fundamental property of the network dynamics.

**Appendix 2 Table 1:** Parameter values for reproducing experimental biases. *σ* values are expressed in fractions of azimuth range in pixels which was set to 360 in our simulations.

| Parameter | Value range | Description |
| --- | --- | --- |
| $K_p$ | 0.4–0.8 | Inhibitory gain |
| $I_0$ | 0.03–0.24 | Prior strength |
| $\sigma_p$ | 0.18–0.25 | Prior width |
| $\sigma_\kappa$ | 0.02–0.12 | Inhibition width |
| $\tau$ | 60 ms | Membrane time constant |
| $N$ | 181 | Ring population size |
| $N_{\text{samples}}$ | 400 | Gaze-centered flow samples |

#### Simulation parameters

The specific parameter values used to produce Figure S16 are: *I*_0_ = 0.03, *K*_*p*_ = 0.4, *σ*_*p*_ = 0.18, *σ*_*k*_ = 0.12.

#### Heading Estimation and 3D Path Reconstruction

The neural heading estimate *θ*_*px*_, decoded from the ring attractor’s peak activity, was converted from image coordinates to an ego-centric heading angle *θ*_*rad*_ using a pinhole camera model. Before converting to angular units, we smoothed the ring attractor’s decoded signal to align with psychophysical evidence that complex motion is integrated over longer time scales than local signals (Burr and Santoro, 2001). We used a temporal window of 2.4 seconds. Given a horizontal field of view (*F OV*_*H*_ = 100^°^) and an image width (*W* = 800 pixels), the camera’s focal length in pixels was defined as:

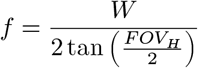

The instantaneous heading in radians was then calculated by taking the inverse tangent of the pixel displacement relative to the principal point:

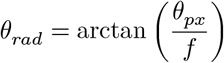

To reconstruct the locomotor path in 3D space, we assumed a constant forward velocity *v* = 0.7 m/s. Under the assumption of a flat ground plane (fixed height *y*), the agent’s position in world coordinates (*X, Z*) was updated via path integration. For each time step Δ*t* = 0.033s, the position transition was defined as:

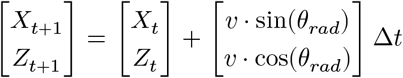

where *Z* represents the forward depth axis and *X* the lateral displacement. This way we were able to compare the reproduced path from the network model with the experimental data.

#### Relation to the controller (phenomenological model)

The neural implementation provides a mechanistic basis for the phenomenological relationship 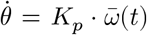 by demonstrating how rotational flow is converted into a change in perceived heading. In the neural model, *K*_*p*_ is not a single fixed parameter, but rather a dynamic property of the recurrent network. The shift in the activity bump (representing 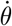) is driven by the spatial asymmetry of the inputs. When the gaze-located inhibition suppresses the sensory evidence at the fixation point, it causes the stable bump of activity to “drift” toward the uninhibited regions of the sensory input. The speed of this drift is directly proportional to the strength of the inhibitory gain (*K*_*p*_ in Equation 28) and the magnitude of the local curl signal (*ω*). Thus, the network naturally performs the integration required by the controller: it transforms localized, gaze-contingent inhibition into a global heading shift that matches the 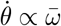 relationship observed in participants.

## Appendix 3

### Controller fits to experimental data

We present the reported perceived headings alongside corresponding model fits derived from different approaches. These approaches vary based on parameter independence:

Separate Fits (Independent Parameters): Parameters are fitted independently for each combination of heading, gaze eccentricity, and retinal flow manipulation. Joint Fits (Common Parameters): A single set of parameters is used across all heading and gaze eccentricity conditions. Each approach was tested with two parameters (the controller gain, *K*_*p*_, and the forward velocity, *v*). The initial motivation for including forward velocity in the fit was to compensate for variations in the timing of perceived heading responses. However, the resulting fitted velocity values (mean ± SD: *v* = 0.87 ± 0.12 m/s) were consistently close to the simulated physical speed during the steady phase (see Fig. 1 Figure Supplement 1).

**Appendix 3 Table 1:** Performance for the two fitting approaches (separate and join: models) in the different retinal flow conditions. Cost is the average 2D deviation per step between the fit and the observed heading.

Model Evaluation Metrics
| Model Type | Flow | Normalized Cost | Negative log-like | AIC |
| --- | --- | --- | --- | --- |
| Separate (30 params) | unaltered | 0.080 | -3.54 | 67.08 |
| Separate (28 params) | cancelled | 0.066 | -3.41 | 62.82 |
| Separate (28 params) | overcancelled | 0.077 | -3.49 | 62.98 |
| Join (2 param) | unaltered | 0.379 | -4.32 | 12.65 |
| Join (2 param) | cancelled | 0.252 | -4.08 | 12.17 |
| Join (2 param) | overcancelled | 0.330 | -4.22 | 12.44 |

- Figures from Figure 4 Fig. Suppl. 2 to Suppl. 7: separate fits
- Figures from Figure 4 Fig. Suppl. 8 to Suppl. 13: joint fits.

### Controller fits to Exp 3 from Wilkie and Wann (2003)

To evaluate the generalizability of the curl-based controller, we fitted the controller to the human steering trajectories reported by Wilkie and Wann (2003). In order to obtain the initial image curl signals by simulating an observer kinematics over an 8-second duration on the same floor of our experiments with initial gaze eccentricities as in Wilkie and Wann (2003) and computed the mean curl using the generated videos of the simulations according to eq. 1. The controller transformed the instantaneous mean curl into a turn rate (*ω*) by adjusting the proportional gain (*K*_*p*_) and the forward velocity (*v*) which minimized the 2D distance between the simulated paths and the steering trajectories by Wilkie and Wann (2003). As shown in S17 in the SI, the model accurately captures the steering behavior across different target eccentricities, with the predicted paths (red) closely aligning with the steering trajectories (grey). Table S3 in the SI shows the fitted parameters and the final lateral position which always ended within the target. The controller gain *K*_*p*_ was very similar in all paths indicating that the differences were caused by the curl signal. The fitted speed was very close to the experimental speed (8 m/s) used in Wilkie and Wann (2003).

**Appendix 3 Fig 1:**
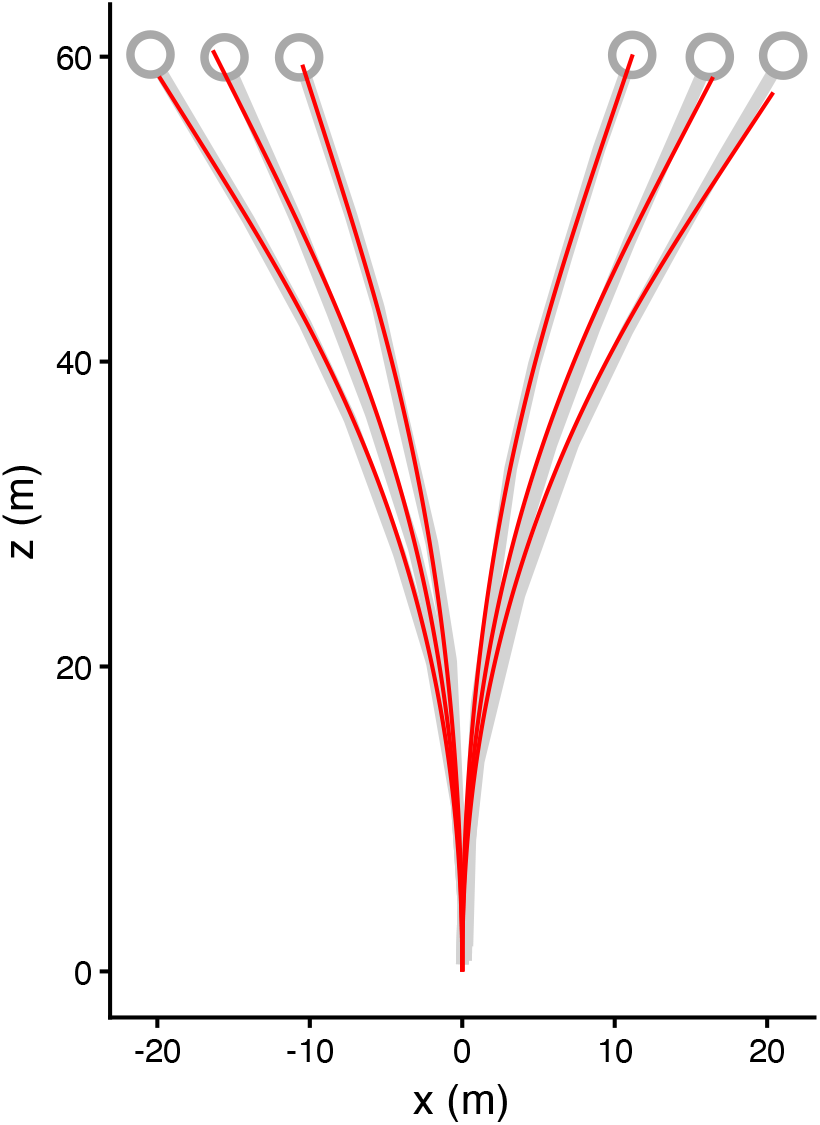
Reproduction of steering paths towards targets. Response of the controller model (red lines) to the conditions introduced in Experiment 3 of Wilkie and Wann 2003. Targets were placed 60 m ahead at 10, 14, and 18 to the left and right (grey circles). The grey lines denote an approximation of the observer paths reported in Wilkie and Wann (2003). The simulated speed in their study was 8 m/s

**Appendix 3 Table 2:** Parameters of the controller to successfully steer to the 6 targets shown in figure S17. The controller speed was very close to the simulated speed (8 m/s) in (Wilkie and Wann, 2003). The final lateral position always ended within the target dimensions (2 m width).

| Target initial eccentricity ($^{\circ}$) | Kp | Vel (m/s) | Final $x$ (m) |
| --- | --- | --- | --- |
| -18$^{\circ}$ | 3.54 | 8.00 | -19.9 |
| -14$^{\circ}$ | 3.57 | 8.03 | -16.4 |
| -10$^{\circ}$ | 3.21 | 7.7 | -10.5 |
| 10$^{\circ}$ | 3.38 | 7.8 | 11.2 |
| 14$^{\circ}$ | 3.69 | 7.82 | 16.4 |
| 18$^{\circ}$ | 3.68 | 7.90 | 20.4 |

