## Supplemental figures for "Beyond the Focus of Expansion: Retinal curl as a functional signal for heading estimation"

##### Supplementary Information

Kontessa I. Zorpala and Joan López-Moliner

#### Video S1

Retinal flow pattern during forward translation with fixation on a target (white dot) located to the left of the heading. For optimal viewing, the display should be centered relative to the observer with a field of view exceeding 50º (e.g., a 60 cm screen viewed from a distance of 60 cm).

#### Video S2

The same as video S1, retinal flow pattern during forward translation and fixating a target (white dot) located to the left of the current path, but, the rotational component of the flow has been counteracted, so there is very little curl around the fovea. For optimal viewing, the display should be centered relative to the observer with a field of view exceeding 50º (e.g., a 60 cm screen viewed from a distance of 60 cm).

#### Video S3

Controller illustration: how heading is corrected to match gaze by applying equation 5 in the main text.

#### Walking speed profile in the simulated display

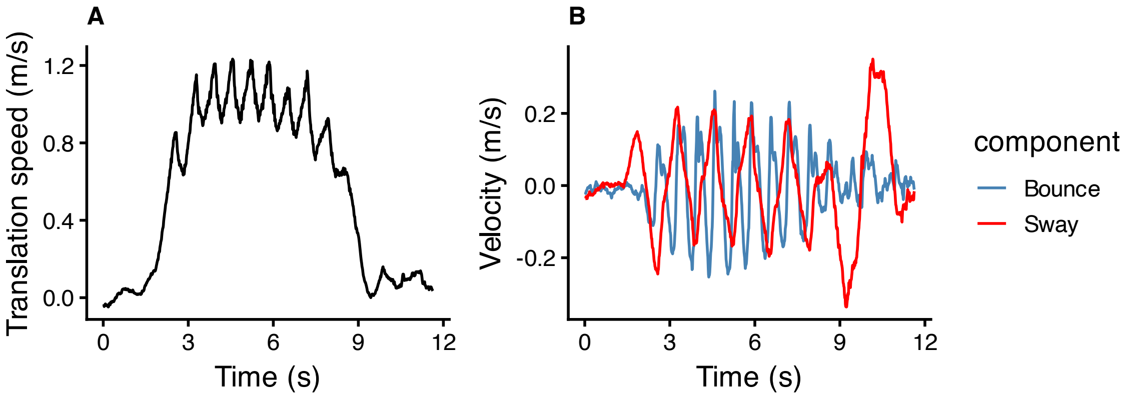


Figure 1 - Suppl. 1: Speed profile for a representative trial for different components. (A) Translational or forward speed. (B) Bounce (vertical component) and Sway (lateral component)

#### Rotation rate

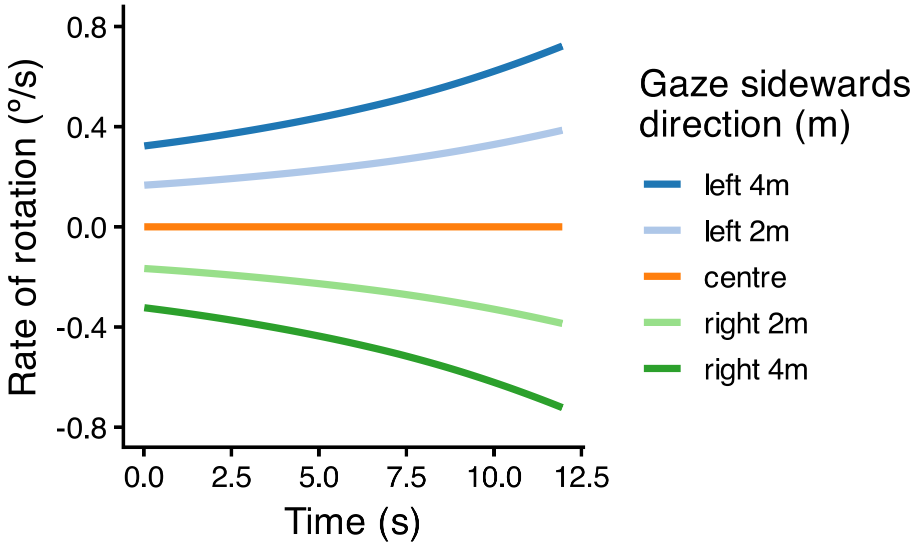


Figure 1 - Suppl. 2: Time course of rotation rate (head/eye)

#### Example of time series raw responses

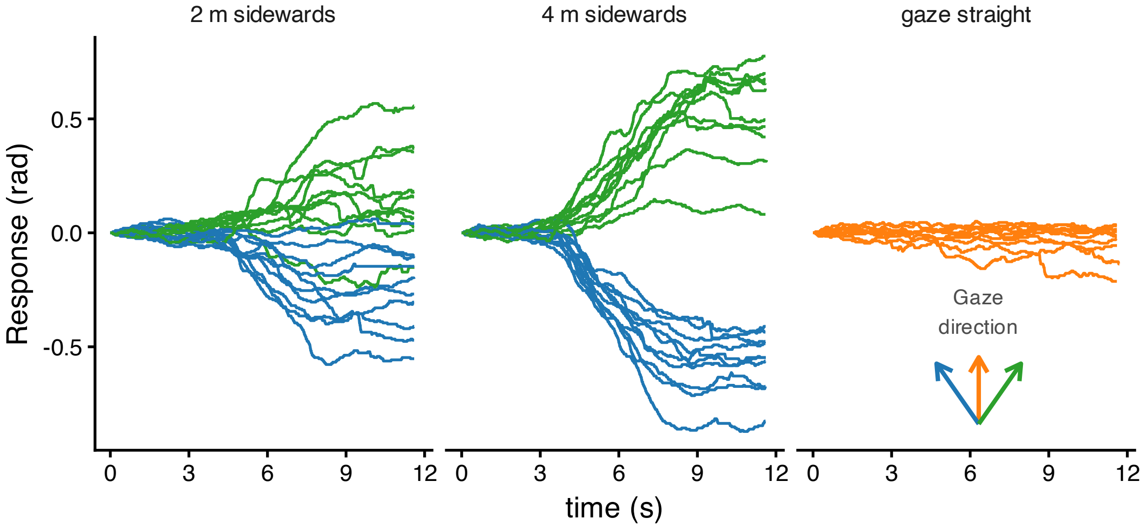


Figure 2 - Suppl. 1: Time course of raw responses for one representative participant in trials simulating straight ahead movement

#### Joint fit to averaged data

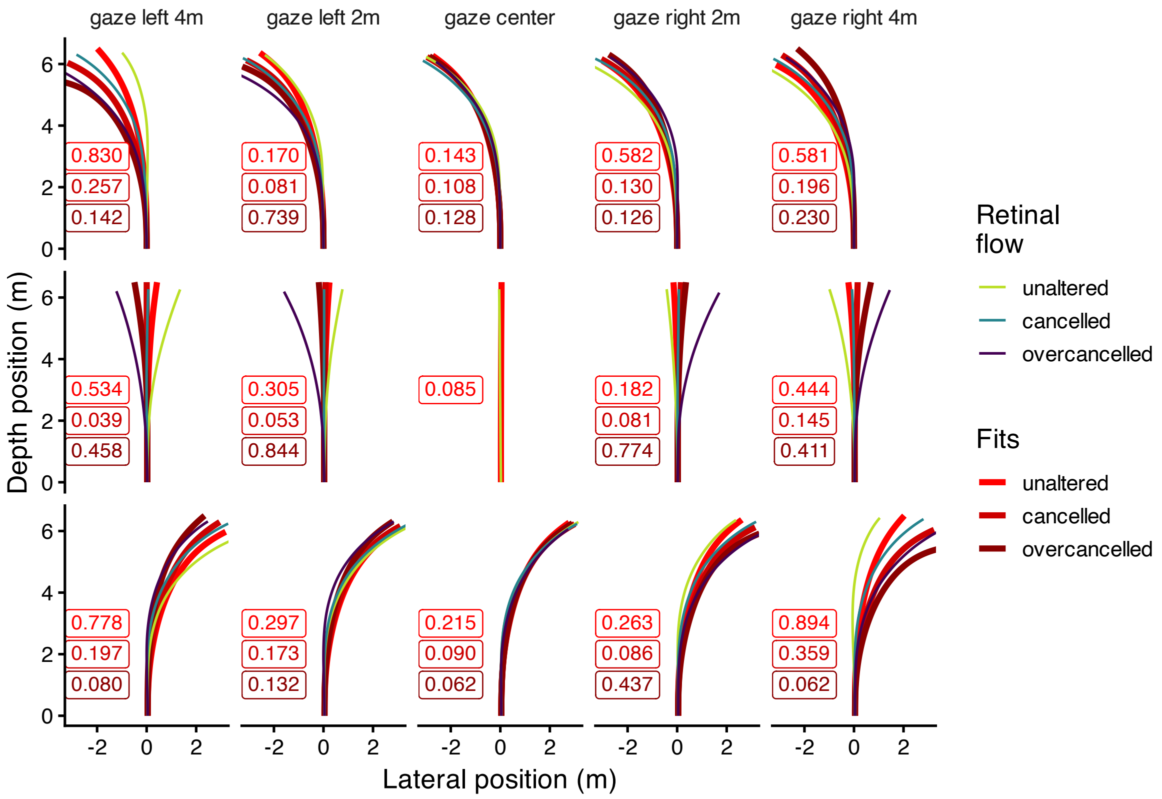


Figure 4 - Suppl. 1: **Joint fits of the controller**. Average perceived heading across participants for each gaze condition (columns) and each physical path curvature (rows), separately for the three retinal–flow manipulation conditions (thiner solid lines): unaltered curl (yellow-green), cancelled curl (cyan), and over-cancelled curl (purple). The different red thicker solid lines denote the best fit of the controller. The numbers in each panel indicate the average lateral deviation per step between the fit and the observed heading. Note that for the centered gaze and straight path, only the unaltered flow condition is shown.

#### Figures with individual fits of the controller

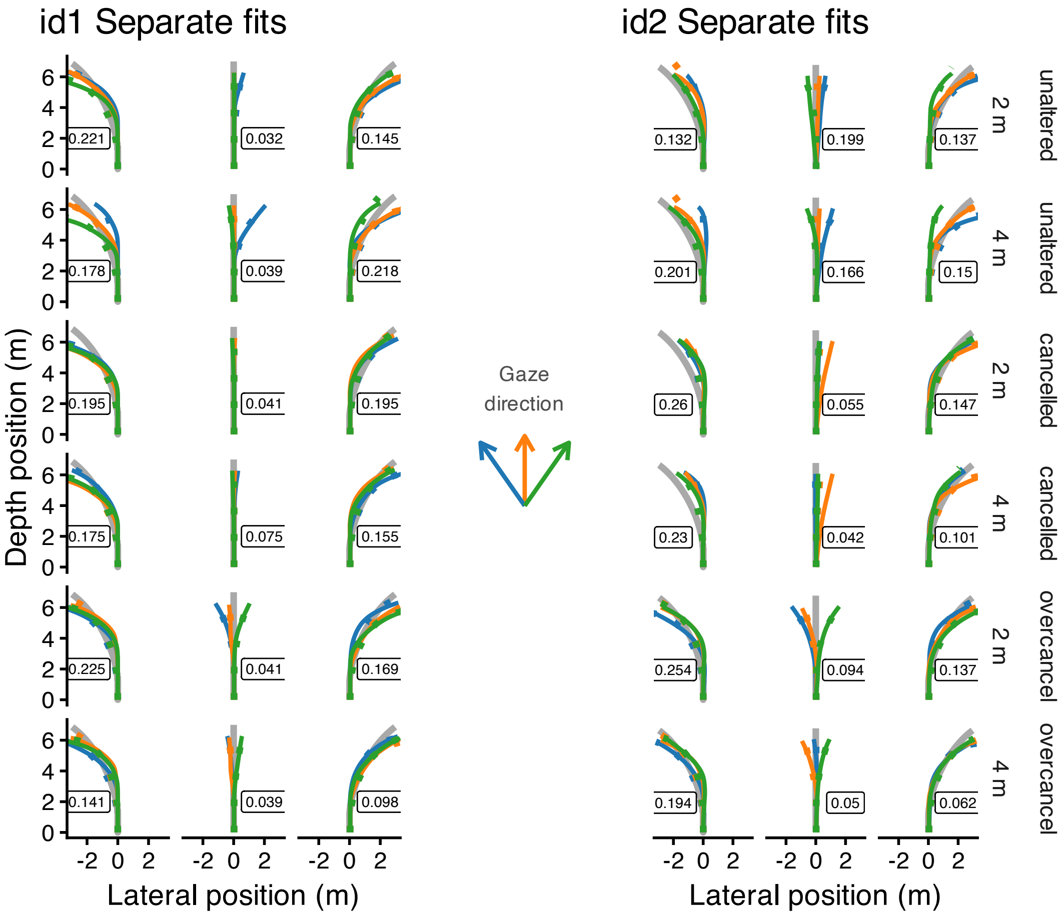


Figure 4 - Suppl. 2: Observed mean heading directions (solid lines) and model fits (dotted lines) are shown for two participants (identified by id#). The plots are organized by heading (columns) and gaze eccentricity (2m/4m) and retinal flow condition (rows within the facets). The color of the trajectories indicates the direction of gaze. The number displayed within each panel represents the mean lateral deviation between the fitted trajectory and the observed path.

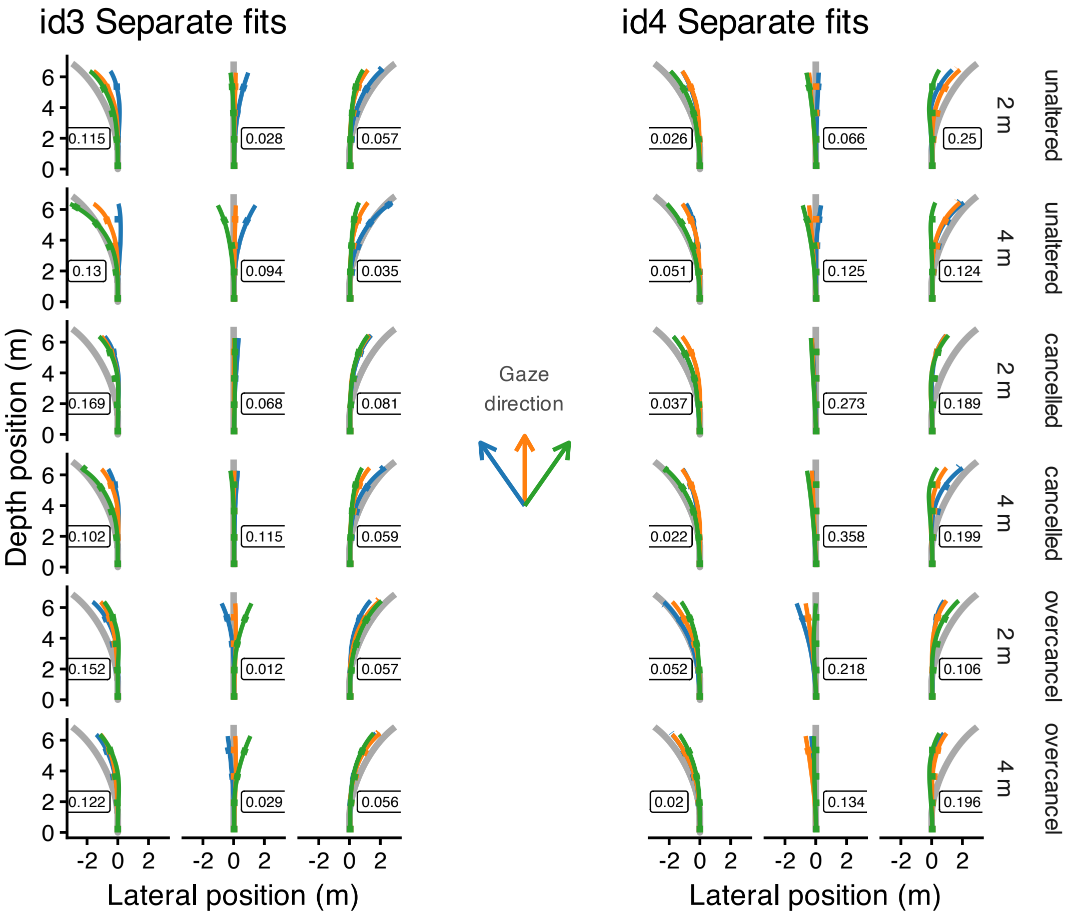


Figure 4 - Suppl. 3: Observed mean heading directions (solid lines) and model fits (dotted lines) are shown for two participants (identified by id#). The plots are organized by heading (columns) and gaze eccentricity (2m/4m) and retinal flow condition (rows within the facets). The color of the trajectories indicates the direction of gaze. The number displayed within each panel represents the mean lateral deviation between the fitted trajectory and the observed path.

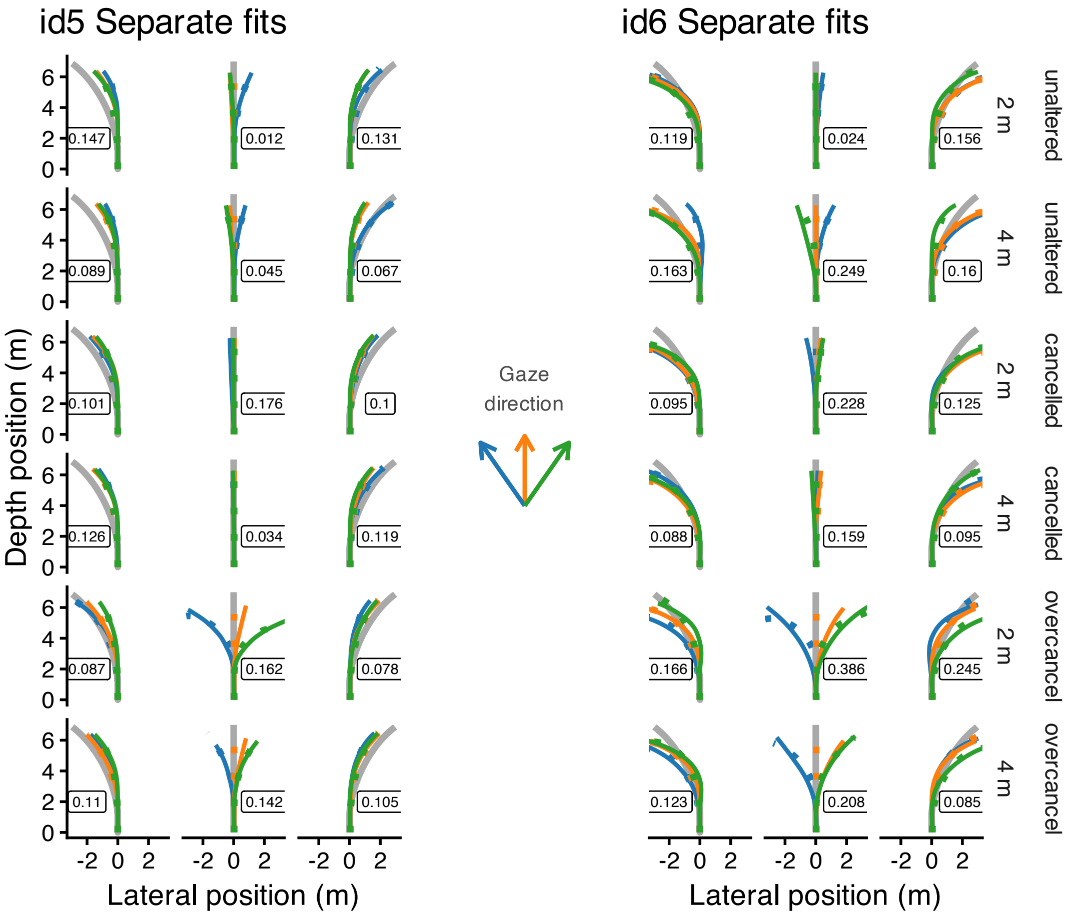


Figure 4 - Suppl. 4: Observed mean heading directions (solid lines) and model fits (dotted lines) are shown for two participants (identified by id#). The plots are organized by heading (columns) and gaze eccentricity (2m/4m) and retinal flow condition (rows within the facets). The color of the trajectories indicates the direction of gaze. The number displayed within each panel represents the mean lateral deviation between the fitted trajectory and the observed path.

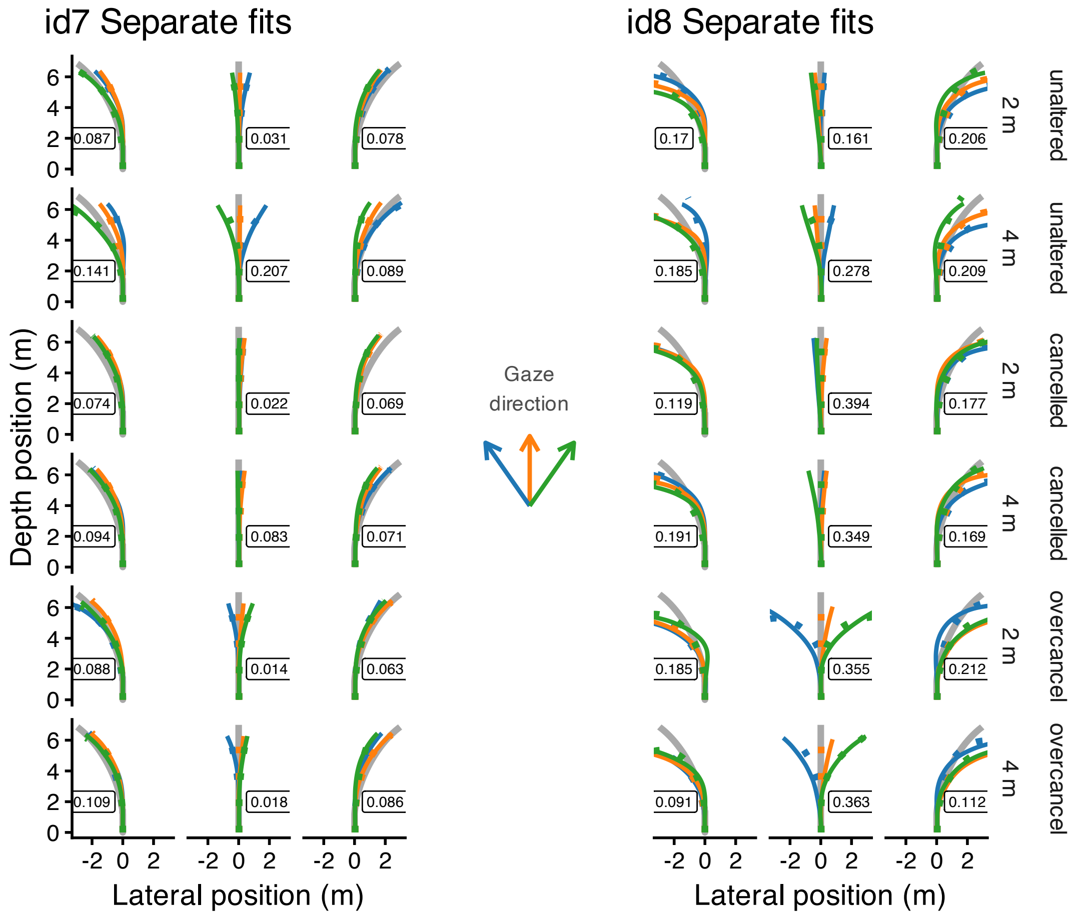


Figure 4 - Suppl. 5: Observed mean heading directions (solid lines) and model fits (dotted lines) are shown for two participants (identified by id#). The plots are organized by heading (columns) and gaze eccentricity (2m/4m) and retinal flow condition (rows within the facets). The color of the trajectories indicates the direction of gaze. The number displayed within each panel represents the mean lateral deviation between the fitted trajectory and the observed path.

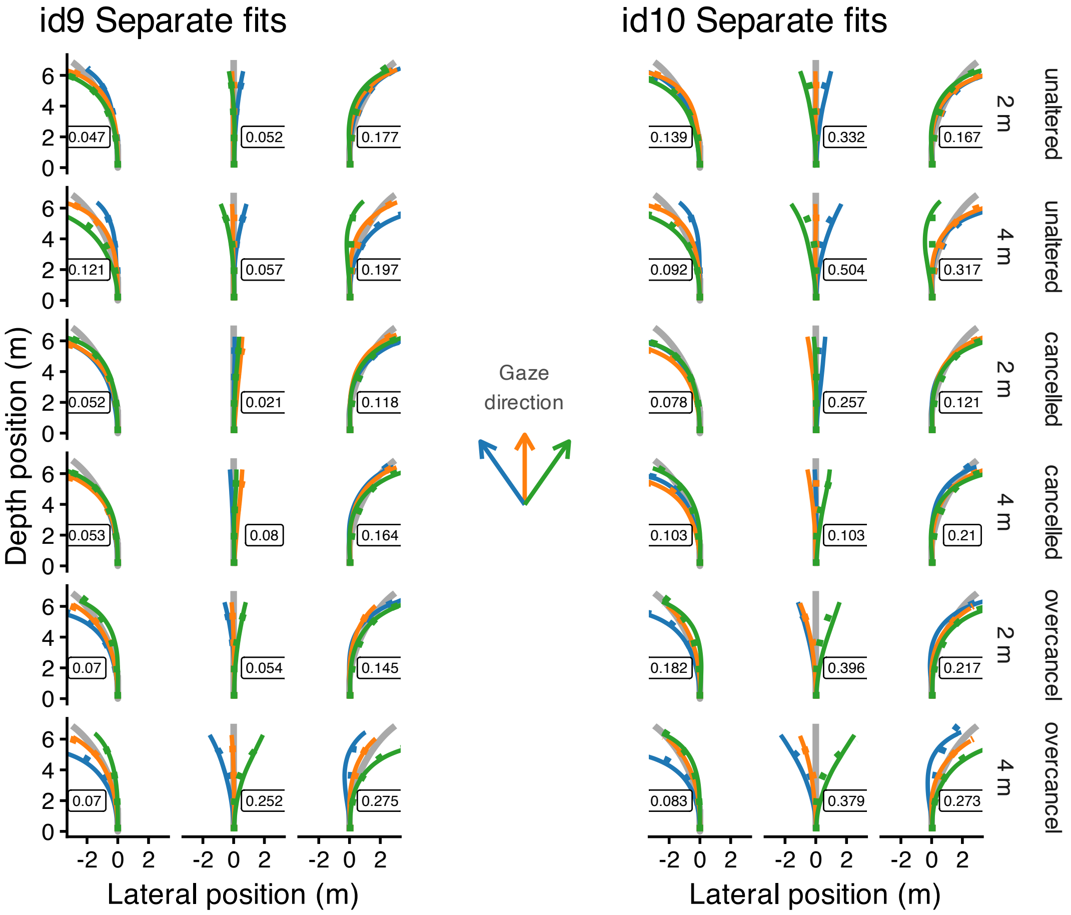


Figure 4 - Suppl. 6: Observed mean heading directions (solid lines) and model fits (dotted lines) are shown for two participants (identified by id#). The plots are organized by heading (columns) and gaze eccentricity (2m/4m) and retinal flow condition (rows within the facets). The color of the trajectories indicates the direction of gaze. The number displayed within each panel represents the mean lateral deviation between the fitted trajectory and the observed path.

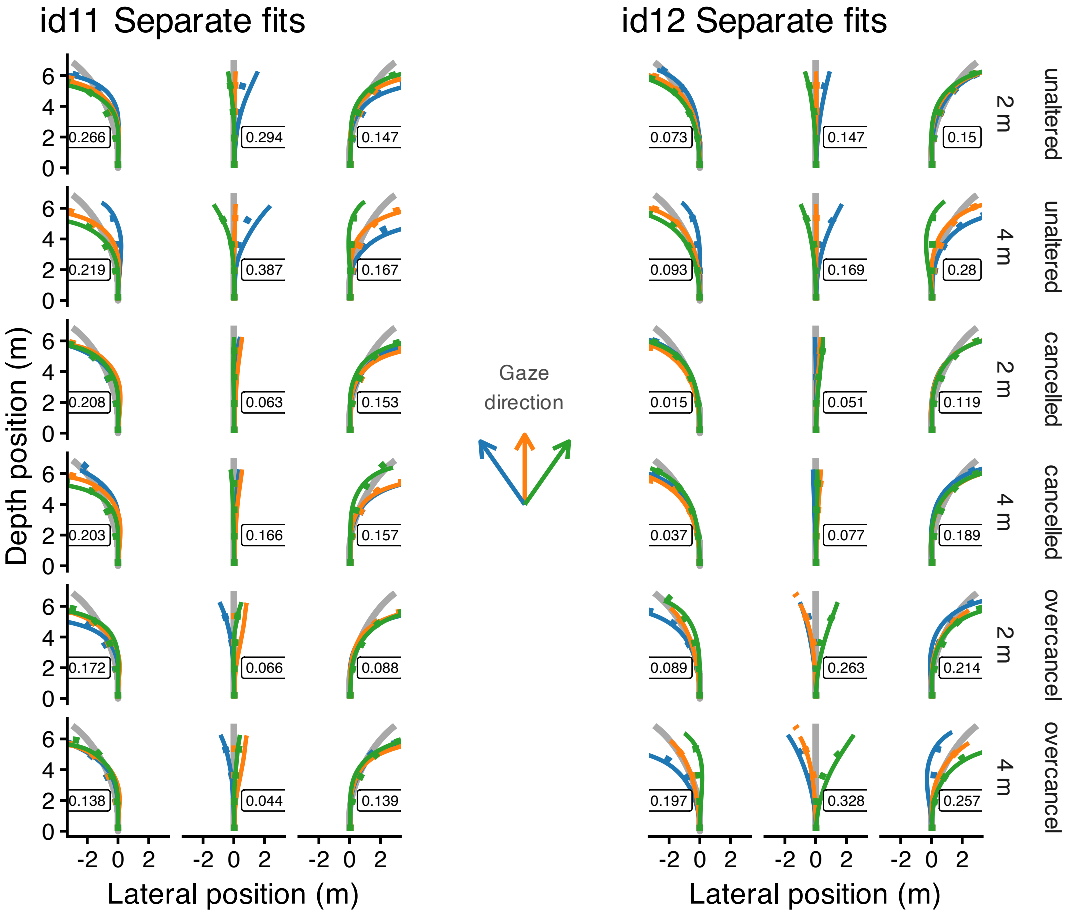


Figure 4 - Suppl. 7: Observed mean heading directions (solid lines) and model fits (dotted lines) are shown for two participants (identified by id#). The plots are organized by heading (columns) and gaze eccentricity (2m/4m) and retinal flow condition (rows within the facets). The color of the trajectories indicates the direction of gaze. The number displayed within each panel represents the mean lateral deviation between the fitted trajectory and the observed path.

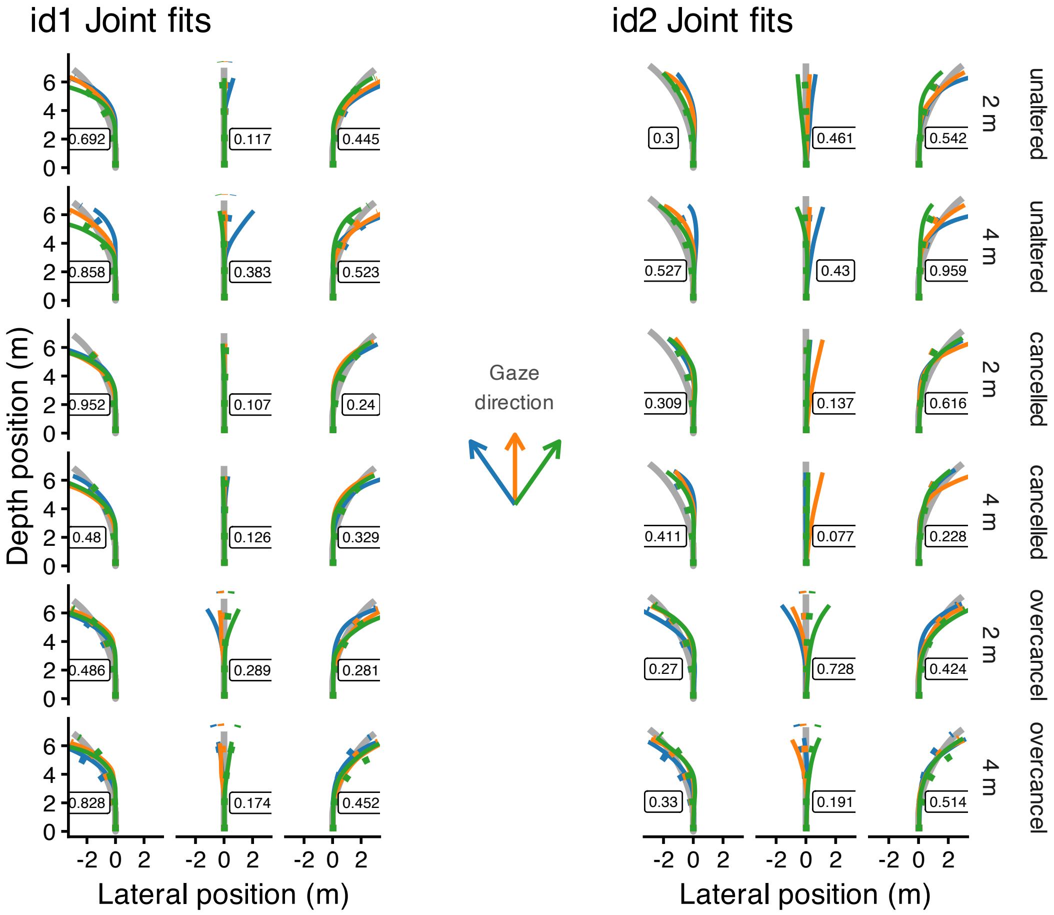


Figure 4 - Suppl. 8: Observed mean heading directions (solid lines) and model fits (dotted lines) are shown for two participants (identified by id#). The plots are organized by heading (columns) and gaze eccentricity (2m/4m) and retinal flow condition (rows within the facets). The color of the trajectories indicates the direction of gaze. The number displayed within each panel represents the mean lateral deviation between the fitted trajectory and the observed path.

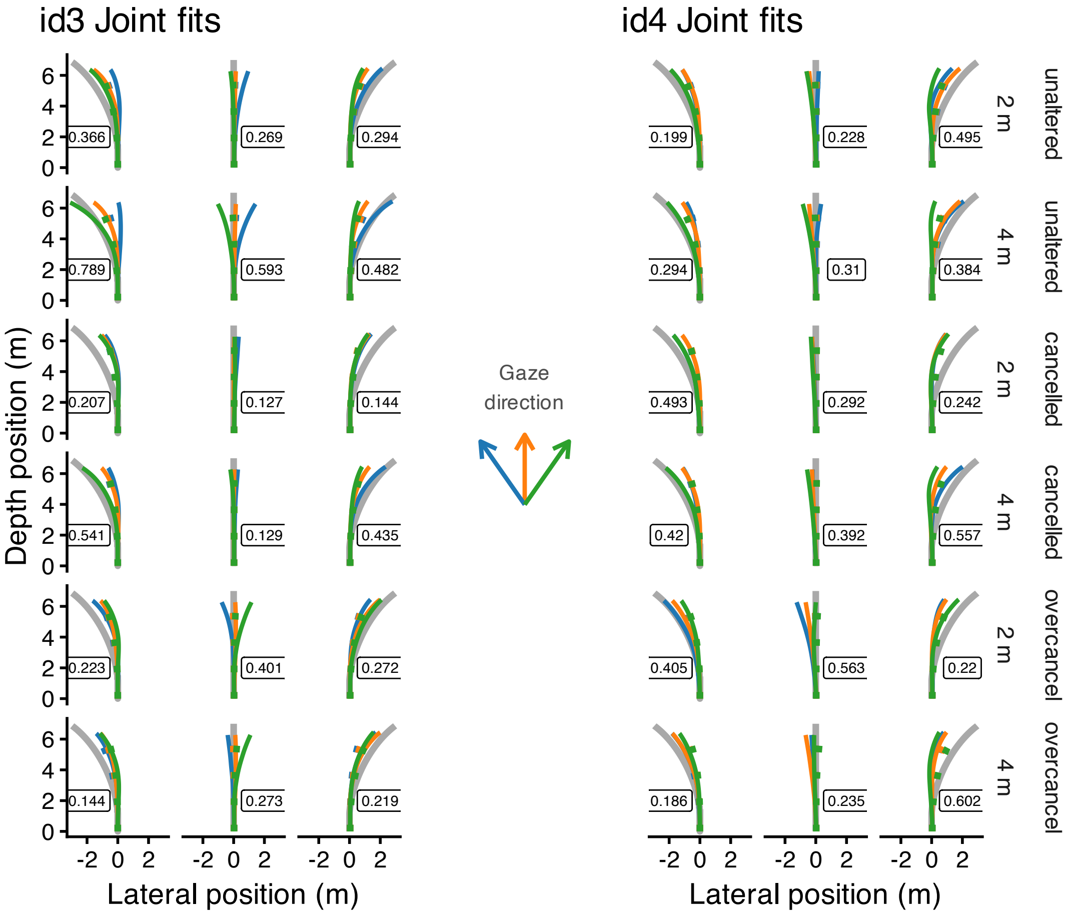


Figure 4 - Suppl. 9: Observed mean heading directions (solid lines) and model fits (dotted lines) are shown for two participants (identified by id#). The plots are organized by heading (columns) and gaze eccentricity (2m/4m) and retinal flow condition (rows within the facets). The color of the trajectories indicates the direction of gaze. The number displayed within each panel represents the mean lateral deviation between the fitted trajectory and the observed path.

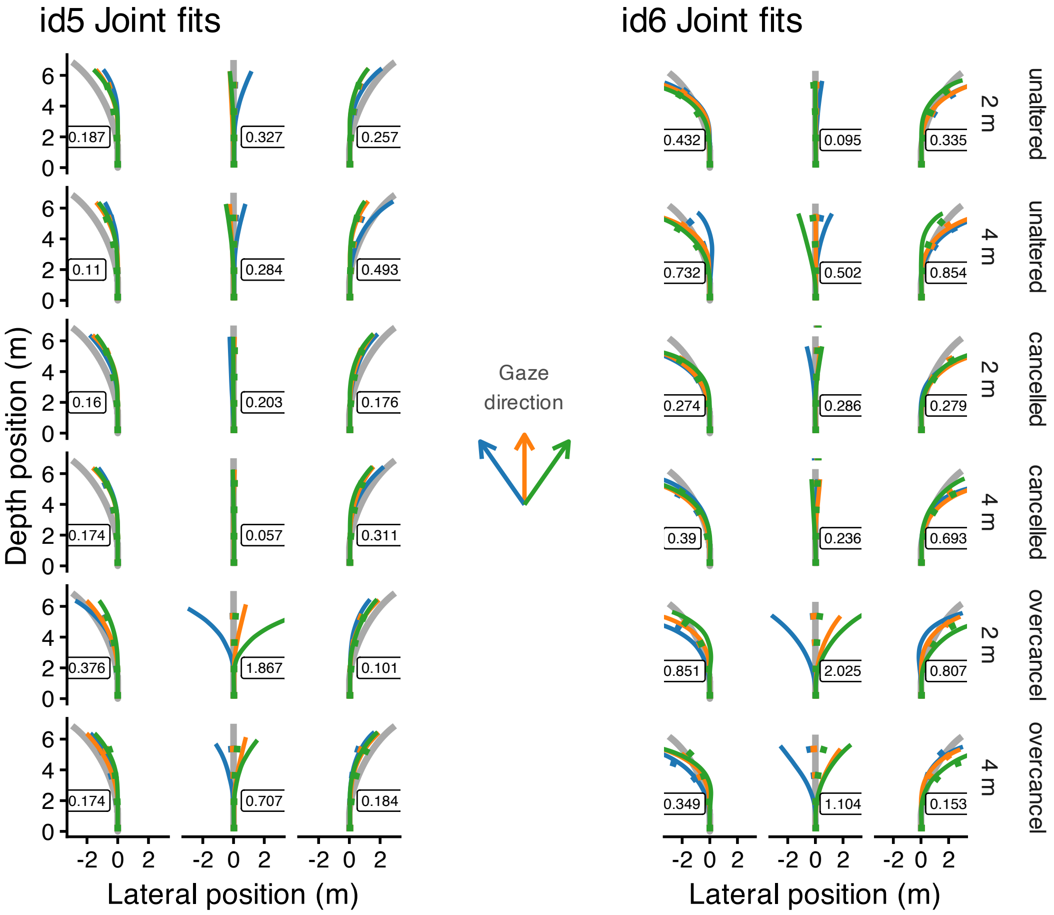


Figure 4 - Suppl. 10: Observed mean heading directions (solid lines) and model fits (dotted lines) are shown for two participants (identified by id#). The plots are organized by heading (columns) and gaze eccentricity (2m/4m) and retinal flow condition (rows within the facets). The color of the trajectories indicates the direction of gaze. The number displayed within each panel represents the mean lateral deviation between the fitted trajectory and the observed path.

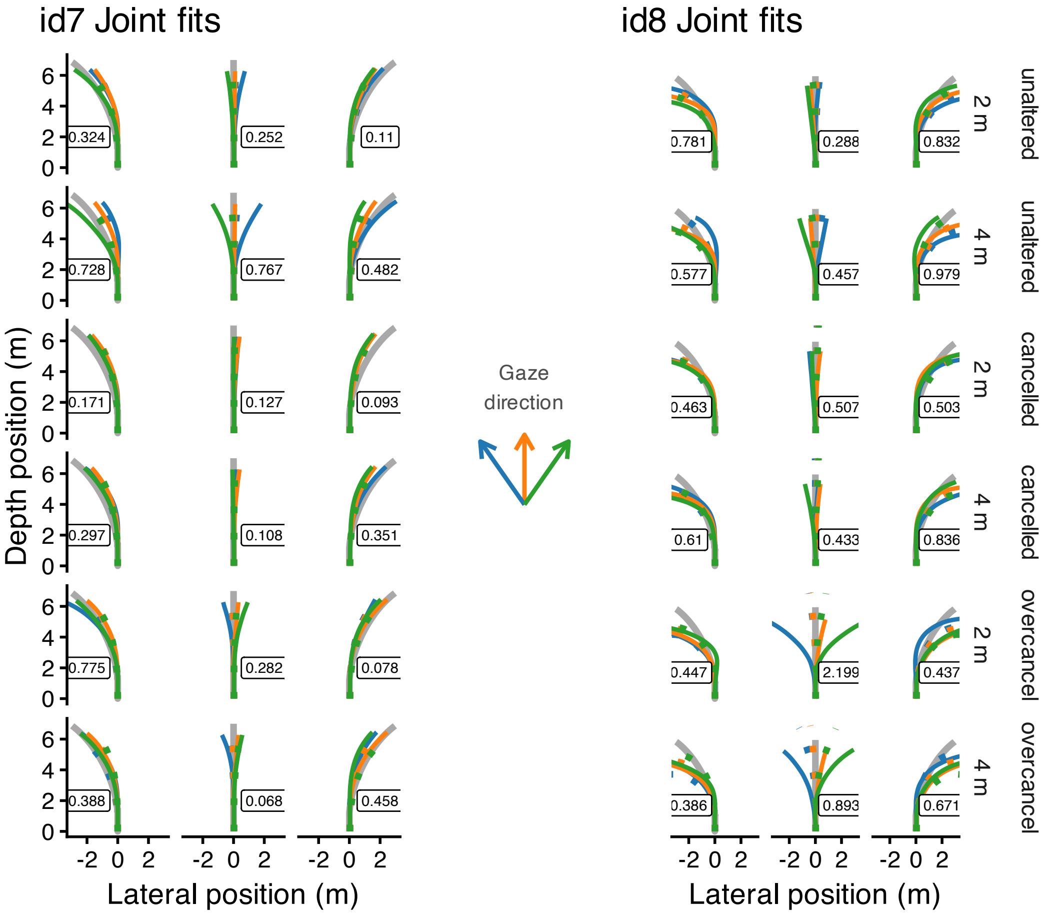


Figure 4 - Suppl. 11: Observed mean heading directions (solid lines) and model fits (dotted lines) are shown for two participants (identified by id#). The plots are organized by heading (columns) and gaze eccentricity (2m/4m) and retinal flow condition (rows within the facets). The color of the trajectories indicates the direction of gaze. The number displayed within each panel represents the mean lateral deviation between the fitted trajectory and the observed path.

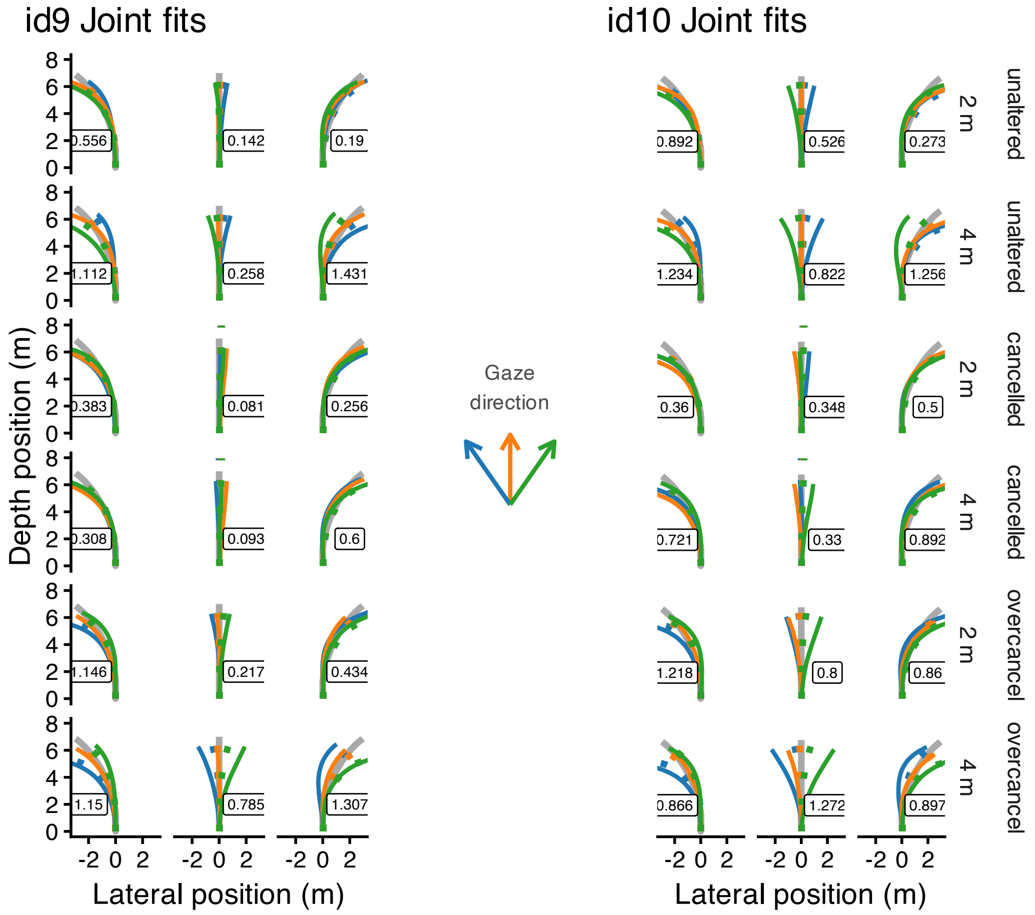


Figure 4 - Suppl. 12: Observed mean heading directions (solid lines) and model fits (dotted lines) are shown for two participants (identified by id#). The plots are organized by heading (columns) and gaze eccentricity (2m/4m) and retinal flow condition (rows within the facets). The color of the trajectories indicates the direction of gaze. The number displayed within each panel represents the mean lateral deviation between the fitted trajectory and the observed path.

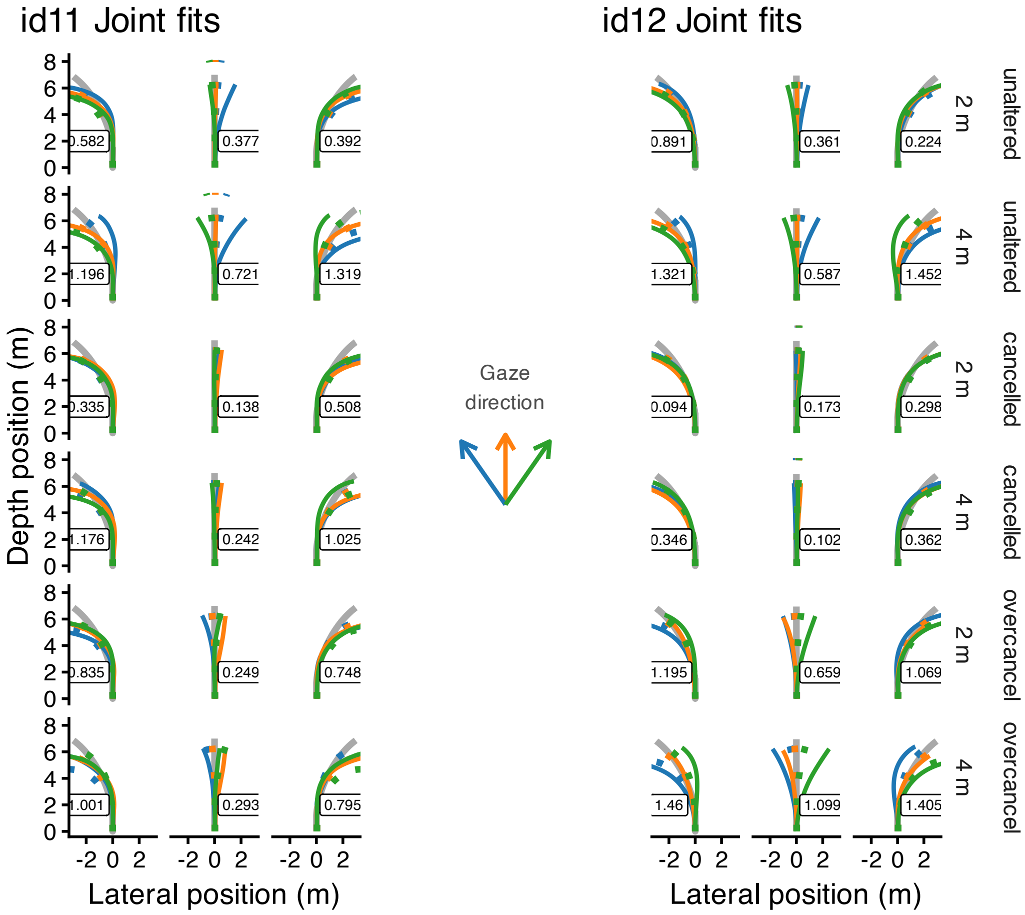


Figure 4 - Suppl. 13: Observed mean heading directions (solid lines) and model fits (dotted lines) are shown for two participants (identified by id#). The plots are organized by heading (columns) and gaze eccentricity (2m/4m) and retinal flow condition (rows within the facets). The color of the trajectories indicates the direction of gaze. The number displayed within each panel represents the mean lateral deviation between the fitted trajectory and the observed path.

### Neural model dynamics

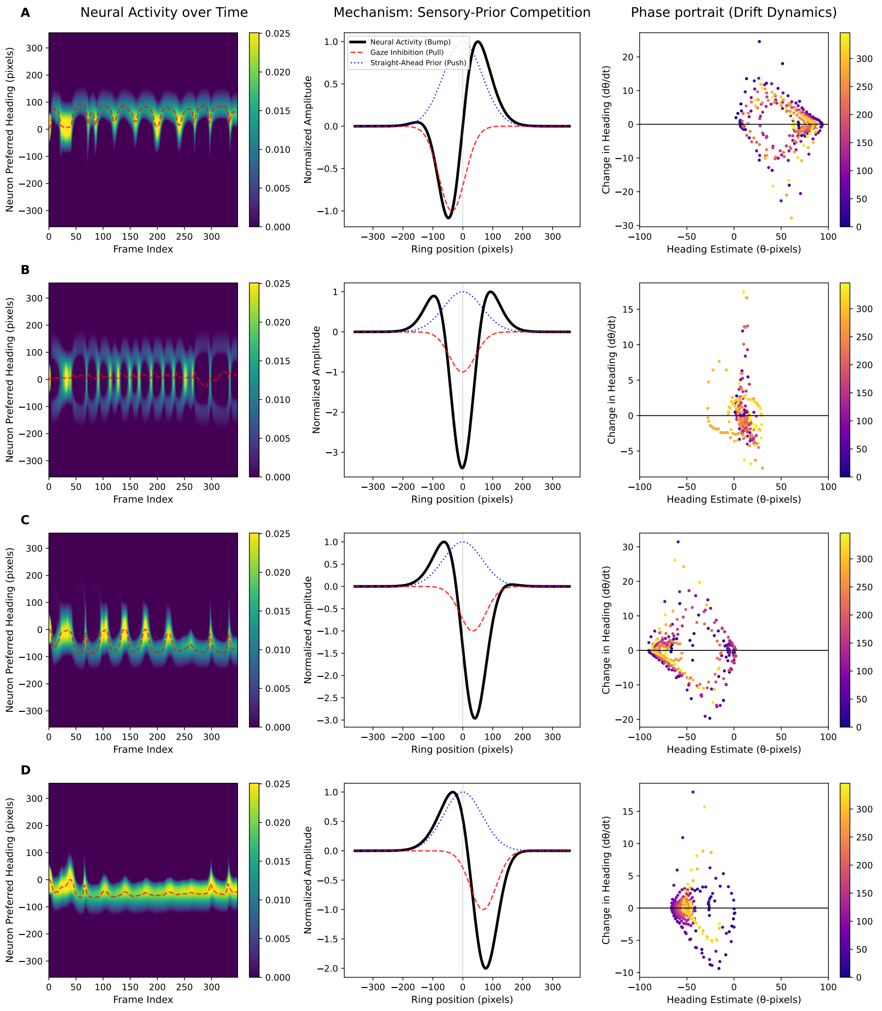


Figure 5 - Suppl. 1: Neural model. (first column) The heatmap shows the neural activity across neuron preferred headings (y-axis) over the frame index (x-axis), with a dashed white line indicating the decoded population heading estimate. The color bar denotes the activity level. (middle column) The mechanism of sensory-prior competition plotting normalized amplitude against ring position in pixels, illustrating the spatial alignment of the straight-ahead prior (blue dotted line), gaze-centered inhibition (red dashed line), and the resulting neural activity bump (black solid line). (last colum) Phase portrait showing the change in heading ($d\theta/dt$) versus the heading estimate ($\theta$) in pixels. The color bar denotes frame number. Different gaze eccentricities are shown row-wise (A) gaze to the left -2 m-, (B) gaze centred, (C) gaze to the right -2 m- and (D) gaze to the right -4 m-.

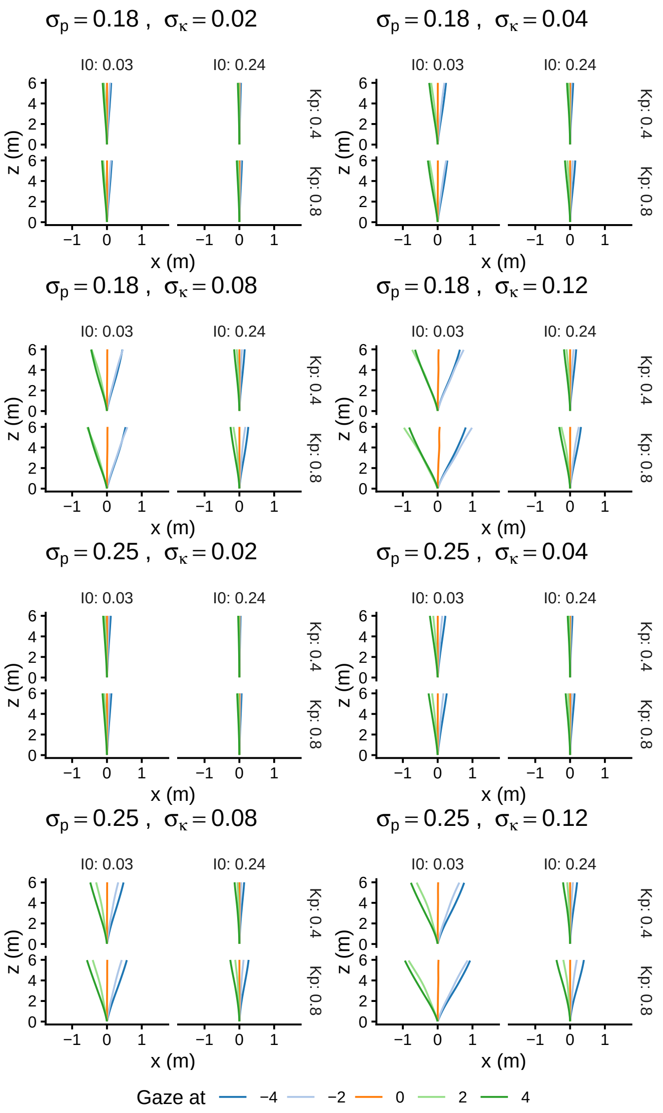


Figure 5 - Suppl. 2: Reproduced 3D paths from the neural model decoding.
